# Integrated histopathology, spatial and single cell transcriptomics resolve cellular drivers of early and late alveolar damage in COVID-19

**DOI:** 10.1101/2023.12.20.572494

**Authors:** Jimmy Tsz Hang Lee, Sam N. Barnett, Kenny Roberts, Helen Ashwin, Luke Milross, Jae-Won Cho, Alik Huseynov, Benjamin Woodhams, Alexander Aivazidis, Tong Li, Joaquim Majo, Patricia Chaves Guerrero, Michael Lee, Antonio M. A. Miranda, Zuzanna Jablonska, Vincenzo Arena, Brian Hanley, Michael Osborn, Virginie Uhlmann, Xiao-Ning Xu, Gary R McLean, Sarah A. Teichmann, Anna M. Randi, Andrew Filby, Paul M. Kaye, Andrew J. Fisher, Martin Hemberg, Michela Noseda, Omer Ali Bayraktar

## Abstract

The most common cause of death due to COVID-19 remains respiratory failure. Yet, our understanding of the precise cellular and molecular changes underlying lung alveolar damage is limited. Here, we integrate single cell transcriptomic data of COVID-19 donor lungs with spatial transcriptomic data stratifying histopathological stages of diffuse alveolar damage (DAD). We identify changes in cellular composition across progressive DAD, including waves of molecularly distinct macrophages and depleted epithelial and endothelial populations throughout different types of tissue damage. Predicted markers of pathological states identify immunoregulatory signatures, including IFN-alpha and metallothionein signatures in early DAD, and fibrosis-related collagens in organised DAD. Furthermore, we predict a fibrinolytic shutdown via endothelial upregulation of *SERPINE1*/PAI-1. Cell-cell interaction analysis revealed macrophage-derived *SPP1*/osteopontin signalling as a key regulator during early DAD. These results provide the first comprehensive, spatially resolved atlas of DAD stages, highlighting the cellular mechanisms underlying pro-inflammatory and pro-fibrotic pathways across alveolar damage progression.

## Main

Since the outbreak of the COVID-19 pandemic in late 2019, SARS-CoV-2 infection continues to spread, with nearly 700 million cases recorded worldwide and almost 7 million deaths (Dong, Du and Gardner, 2020). Whilst being a respiratory illness, the severity across infected patients is variable, with critical cases manifesting as a systemic disease with hyperinflammation and cytokine storm, leading to multiple organ damage and dysfunction. Crucially, endothelial damage and associated coagulopathy are reported as contributing to severe forms of the disease (Varga *et al.*, 2020). A better understanding of the cellular and molecular mechanisms underlying the devastating lung alveolar damage in COVID-19 could inform novel therapies to the benefit of patients with severe symptoms.

The predominant histological lung injury pattern in COVID-19 is termed diffuse alveolar damage (DAD). DAD presents with heterogeneous histopathological features and stages, where early or exudative DAD (EDAD) is characterised by hyaline membrane deposition and inflammation, while late or organising DAD (ODAD) is marked by extensive fibrosis, with intermediate states showing mixed pathological features (MDAD) (Milross, Majo, Cooper, *et al.*, 2022). Hence, EDAD and ODAD display increasingly severe patterns of lung injury, and are thought to represent temporal progression of disease pathology (Cardinal-Fernández *et al.*, 2017; Erjefält *et al.*, 2022; Ashwin *et al.*, 2023; Milross *et al.*, 2023). However, beyond initial reports documenting expanded immune cells and fibroblast populations in ODAD (Erjefält *et al.*, 2022), we lack an unbiased, fine-grained examination of cellular and molecular differences across DAD stages. Furthermore, distinct DAD stages can be spatially intermixed in a given donor’s lung samples (Ashwin *et al.*, 2023; Milross *et al.*, 2023), obfuscating their molecular signatures in tissue dissociation based single cell and bulk RNA-sequencing datasets.

Previously, a wide range of assays, including single cell/single nuclei RNA sequencing (sc/snRNA-seq), spatial transcriptomics (ST) and imaging mass cytometry were successfully applied to study bronchoalveolar fluid and post-mortem lung tissue samples from COVID-19 patients (Bharat *et al.*, 2020; Delorey *et al.*, 2021; Melms *et al.*, 2021). Moreover, targeted profiling using subsets of genes and proteins allowed to define early immune cell recruitment and inflammatory pathway activation, followed by fibrosis, but did not provide a comprehensive overview of the cellular and molecular changes (Delorey *et al.*, 2021; Rendeiro *et al.*, 2021). Consequently, the exact drivers of tissue remodelling within DAD stages remain incompletely understood.

Here, we combine sc/snRNA-seq and ST to provide the first comprehensive cellular and molecular characterisation of DAD stages in COVID-19. We integrated 11 datasets to create a large multi-study sc/snRNA-seq atlas, with a newly generated ST dataset profiling histologically defined DAD stages across autopsy lung tissue samples from 33 patients. For each DAD stage, we identified distinct molecular biomarkers, pathological cell states, tissue microenvironments, as well as cell-cell interactions (Fig. 1A). In particular, we identify waves of macrophage subtypes accumulating through progressive DAD and the enrichment of COVID-19 specific *SPP1* (encoding osteopontin - OPN) signalling from macrophage subpopulations in early DAD. Furthermore, we link *SPP1*/OPN signalling to fibrinolytic shutdown via endothelial upregulation of *SERPINE1* upon OPN treatment. By combining multimodal transcriptomics data and integrating histopathological definition of tissue damage, our study provides a framework that can be applied to other organs in health and disease.

**Figure 1.**
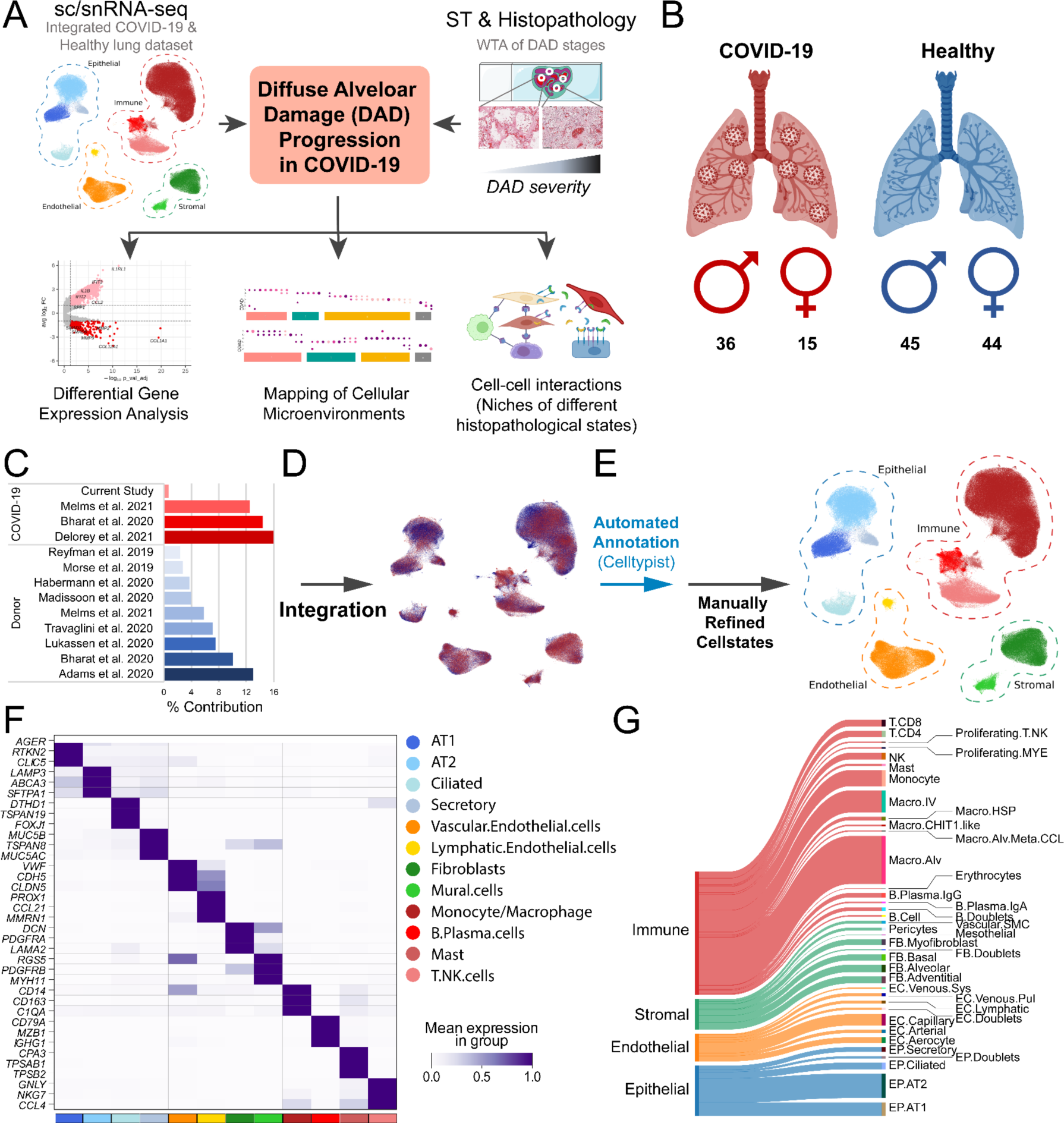
Single cell transcriptomic atlas of the healthy and COVID-19 lung. **A)** Schematic overview. A sc/snRNA-seq dataset was integrated with histopathology driven spatial whole transcriptome analysis. The cell2location tool was used to map cell types/states to spatial transcriptomic data, with DGE, cell-colocalization, and abundance analysis, as well as cell-cell interaction interrogation performed downstream. **B)** Number of COVID-19 patients / donors contributing to sc/snRNA-seq dataset. **C)** Percentage contribution of datasets from organ donors and COVID-19 patients. **D)** UMAP representation of integrated COVID-19 (red) and healthy control (blue) datasets contributing to the final sc/snRNA-seq object. **E)** UMAP representation of the global object with broad cell type (dotted) and mid-level annotation. **F)** Heatmap representation of markers used for mid-level annotation. **G)** Sankey plot visualisation of cell state level annotations derived from subclustering of the broad cell type compartments (Extended Fig. 1*C-D*, Table S3-4).

## Results

### Single cell transcriptomic atlas of the healthy and COVID-19 lung

To maximise the number of COVID-19 lung cells and individuals studied, we generated an integrated sc/snRNA-seq atlas comprising newly generated snRNA-seq data and 10 publicly available sc/snRNA-seq datasets (Fig. 1B-C, Table S1-2). Lung tissue from organ donors (n=89) were used as control in comparison with COVID-19 patients (n=51). After sample processing and quality control, the resulting integrated object comprised 514,756 cells and nuclei (Fig. 1D-E). Integration of transcriptomic data was performed accounting for variations from donor, cell/nuclei, and 10x Genomics chemistry (Fig. 1D, Extended Fig. 1A-C). Leiden clustering was performed and we identified 12 coarse-grained cell states within four major cellular compartments based on curated lineage-specific gene markers and unbiased differential gene expression (DGE) analysis (Fig. 1F). This included epithelial (EP), endothelial (EC), stromal and immune cells, including myeloid (MYE) and lymphoid lineages (Fig. 1E).

Further subclustering defined 32 distinct cell states (Fig. 1G, Extended Figure 1C-D, Table S3-4), including four EP states and six EC states. The latter includes the recently defined EC.Aerocyte (*HPGD*, *EDNRB*, *IL1RL1*) and two *ACKR1*+ venous EC populations (systemic and pulmonary - EC.Venous.Sys, EC.Venous.Pul) distinguished by expression or lack of *COL15A1,* respectively (Gillich *et al.*, 2020; Schupp *et al.*, 2021) (Fig. 1F, Extended Fig. 1D). Vascular smooth muscle cells (Vascular.SMC), pericytes and mesothelial cells were also profiled, along with four fibroblasts (FB) populations within the stromal compartment (Fig. 1F, Extended Fig. 1D). Within immune cells, we identify B plasma cells expressing either *IGHA1* or *IGHG1* (B.Plasma.IgA, B.Plasma.IgG), and five macrophage populations, including an alveolar macrophage subset co-expressing markers from ‘macro-alv-MT’ (including metallothionein (MT) related *MT1F* and *MT1H*) and ‘macro-alv-CCL’ (including chemokines *CCL4* and *CCL20*) identified in previous studies (Madissoon *et al.*, 2022), here termed Macro.Alv.Meta.CCL. (Fig. 1F, Extended Fig. 1D). Additionally, we profile four lymphoid populations, in addition to two proliferating immune populations (Proliferating.T.NK, Proliferating.MYE). Consistent with previous reports (Delorey *et al.*, 2021), we observed widespread gene expression differences in these cell states between healthy and COVID-19 lungs including EP.AT1 and EP.AT2 cell states that showed dysregulation of genes associated with interferon response, and as described below, endothelial cell states that showed dysregulation of the coagulation cascade, (Extended Fig. 1E-G, Table S5-7).

### Transcriptome-wide spatial atlas of DAD stages

Given that DAD stages can be spatially intermixed in COVID-19 lung tissue (Ashwin *et al.*, 2023; Milross *et al.*, 2023), we used ST to identify transcriptional differences across histopathologically defined DAD patterns. For this purpose, we examined post-mortem lung tissue samples from a multi-centre patient cohort that we recently characterised using targeted antibody panels (Ashwin *et al.*, 2023; Milross *et al.*, 2023) Our cohort included 33 individuals, mostly males, across different ages (22-98 years; median 27 years), who died from severe COVID-19 during the first and second wave of the pandemic and spanned diverse ethnic backgrounds and clinical histories, including both hospital (70% of patients) and community (30%) deaths (Extended Fig. 2, Table S8).

**Figure 2.**
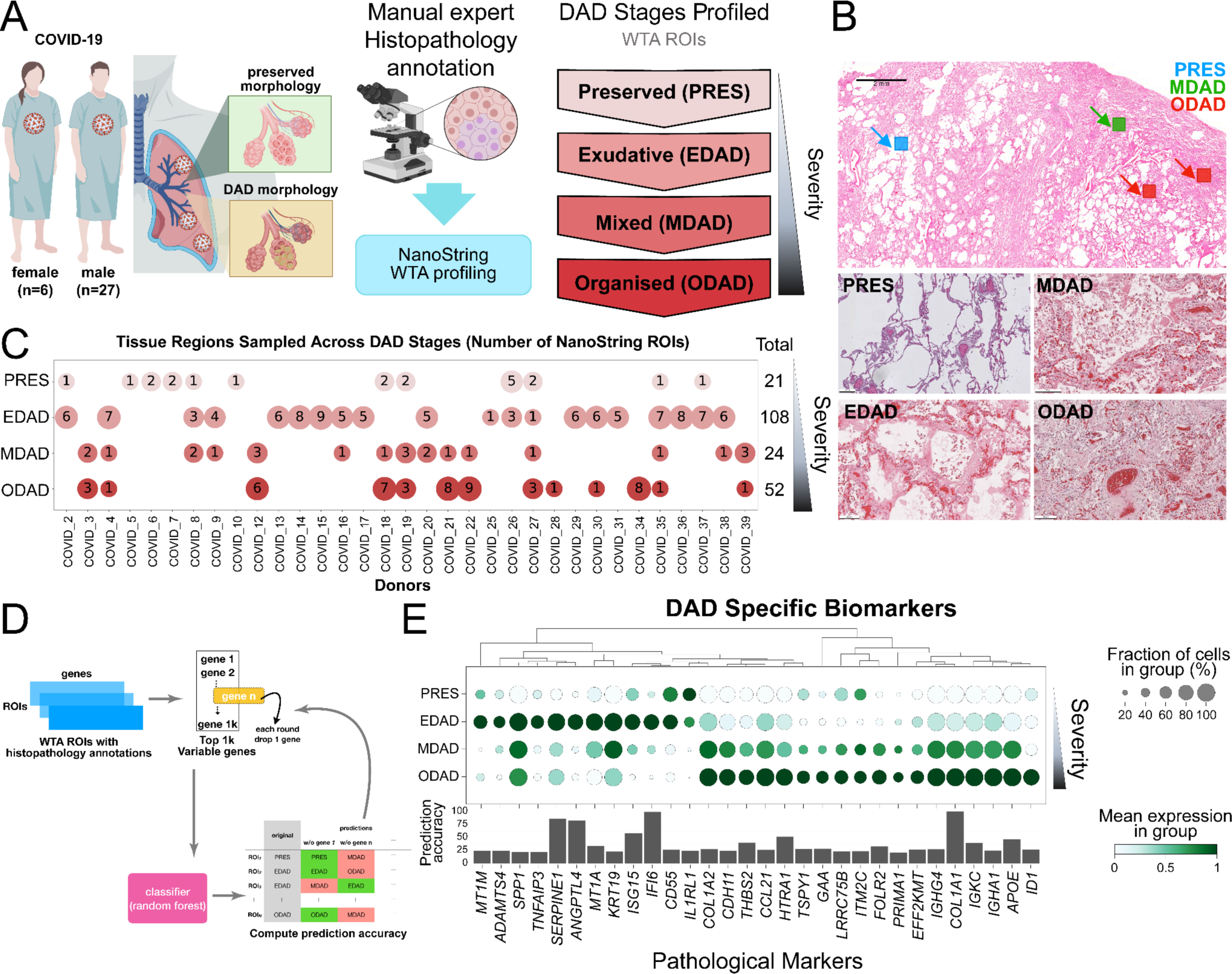
Transcriptome-wide spatial profiling of progressive DAD stages. **A)** Number of COVID-19 patients contributing to spatial WTA dataset (left) and histopathological regions of interest (ROI). **B)** Representative images from H&E stained tissue sections illustrating preserved (PRES) tissue structure and ROIs including different DAD histopathological features. Upper: Indication of multiple ROIs taken from the same tissue section. Lower: Tissue morphology of PRES, EDAD, MDAD and ODAD states. **C)** Number of histopathological ROIs for each COVID-19 patient used in study. **D)** Schematic representing random forest classifier approach for predicting pathology associated gene signatures. **E)** Dotplot representation of histopathological state associated gene signatures obtained from random forest classifier analysis. Dot colour = scaled mean gene expression within a group. Dot size = percentage expressed in group. Prediction accuracy = ratio of correct pathology classification in all ROIs.

Here, we performed unbiased ST characterisation of DAD stages across our patient cohort using the Nanostring Whole Transcriptome assay (WTA), profiling a total number of 326 tissue regions of interest (ROIs) sized 400 µm^2^ each (Fig. 2A). As before, we defined DAD stages strictly based on published histological criteria (Katzenstein, Bloor and Leibow, 1976) and guided by expert pulmonary pathologists (Ashwin *et al.*, 2023; Milross *et al.*, 2023) (Fig. 2B, Methods). We profiled tissue regions with EDAD, MDAD and ODAD as well as tissue areas with preserved (PRES) lung morphology contrasting DAD. To identify the molecular and cellular signatures specific to DADs, we also examined COVID-19 patients with pulmonary oedema consistent with acute cardiac failure as well as superimposed bacterial bronchopneumonia. Finally, as healthy controls, we also sampled lung sections of two patients who died from non-COVID-19 disease and included a publicly available lung WTA dataset of three non-COVID-19 patients (Delorey *et al.*, 2021) in our analysis. In addition, we conducted Visium profiling on six FFPE lung tissue sections, obtained from three patients, to validate the findings of our WTA analysis.

We applied the standard WTA data processing workflow, using stringent gene filters (>6,000 genes per ROI) for quality control and background correction using the CountCorrect algorithm that leverages negative WTA control probes (Roberts *et al.*, 2021). This resulted in 260 high quality spatial transcriptome profiles with an average of 11,423 genes and 689 nuclei detected per ROI (Extended Fig. 3). Our final dataset captures transcriptomic profiles of each DAD stage across multiple patients (Fig. 2C), where multiple DAD stages can often be observed in a given patient (Fig. 2B-C). Hence, our dataset enables a robust comparison of DAD stages accounting for both patient variability and technical batch effects, providing the most comprehensive transcriptomic profiles of DAD progression to date. Our ST data along with our sc/snRNA-seq atlas can be accessed at https://covid19-multiomicatlas.cellgeni.sanger.ac.uk/.

**Figure 3.**
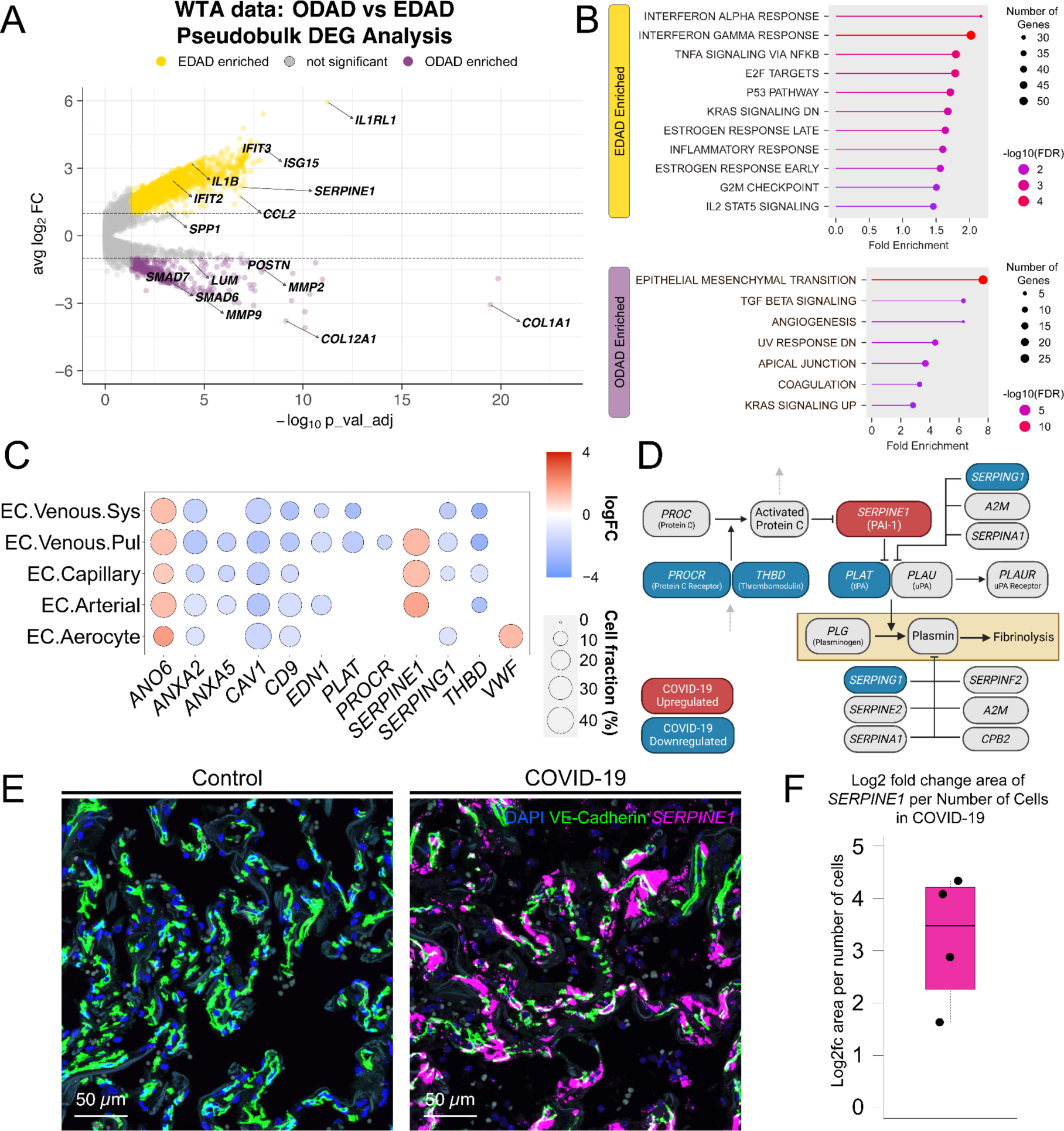
Transcriptional progression of exudative to organised DAD pathology. **A)** Volcano plot representing genes upregulated in EDAD (yellow) and ODAD (purple). **B)** Hallmark MSigDB gene set enrichment of genes upregulated in EDAD (top) vs. ODAD (bottom). **C)** Dot plot illustrates DGE between COVID-19 and controls related to fibrinolysis and coagulation in the sc/snRNA-seq dataset. Red = upregulated in COVID-19, blue = downregulated in COVID-19. For a complete list of DGE, see Table S5. **D)** Schematic representation of the fibrinolysis pathway. Genes upregulated (red) and downregulated (blue) in COVID-19 EC compared to healthy controls are shown. **E)** Healthy control and COVID-19 lung parenchyma samples stained by smFISH for SERPINE1 (magenta), and CDH5 protein (green) counterstained with DAPI (blue). **F)** Boxplot shows log2fc area of SERPINE1 staining per number of cells in COVID-19 lung parenchyma (n=4) compared to healthy controls (n=2).

### Distinct transcriptional signatures of DAD stages

As a first step to determine molecular signatures associated with different DAD states, we used a random forest classifier for the identification of predictive biomarkers of PRES, EDAD, MDAD, and ODAD (Fig. 2D, Methods). Within EDAD ROIs, the classifier identified genes associated with inflammatory response (*SPP1*, *IFI6*, *ISG15*, and *TNFAIP3*), regulation of fibrinolysis (*SERPINE1*), and metallothionein-related genes (*MT1A* and *MT1M*) (Fig. 2E). Whilst metallothionein (MT) genes are typically attributed to metal ion homeostasis and oxidative stress alleviation, they are also involved in early-stage inflammatory responses (Dai *et al.*, 2021), concordant with the pathophysiological phenotype of EDAD in acute lung injury. Further, EDAD is enriched for *ADAMTS4* (aggrecanase-1), a protease upregulated in severe influenza infection, which disrupts lung tissue integrity to enable early immune infiltration by degrading extracellular matrix proteins, including versican (Boyd *et al.*, 2020). Conversely, in ODAD, we observed markers associated with fibrosis, and TGF-beta pathway (*COL1A1*, *COL1A2*, *ID1*, *HTRA1*), and anti-inflammatory response (*FOLR2*) (Puig-Kröger *et al.*, 2009) (Fig 2E). These predicted biomarkers identified from WTA data displayed consistent expression patterns across Visium spots annotated as PRES, EDAD, and ODAD histopathological states (Extended Fig. 4A). Additionally, the top 100 biomarkers exhibited a high correlation between WTA and Visium datasets, with Spearman’s correlation coefficients of 0.56 (p-value=1.4e-9) and 0.52 (p-value=2.6e-8) in EDAD and ODAD, respectively. (Extended Fig. 4B).

**Figure 4.**
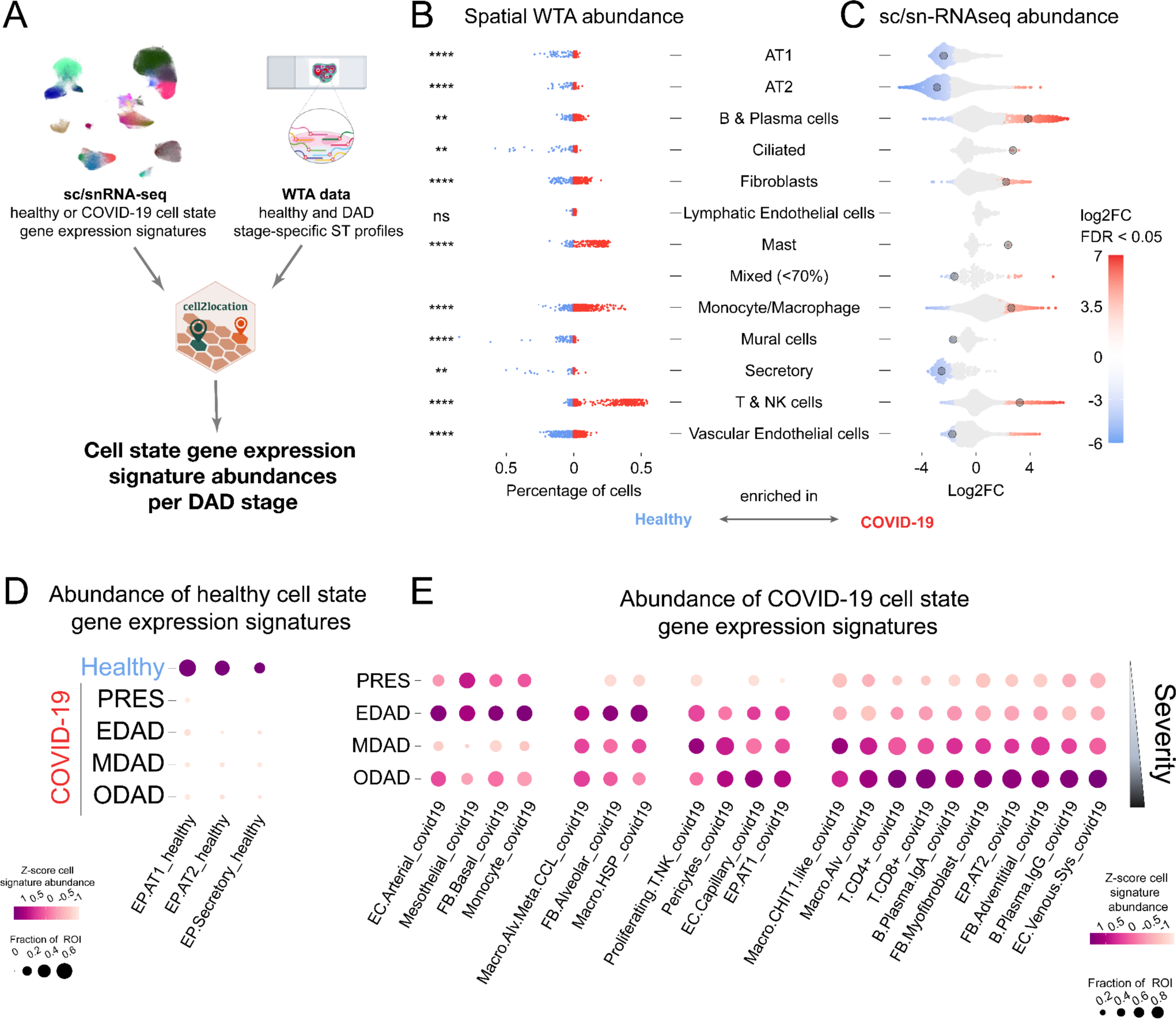
Evolving cellular composition of DAD stages. **A)** Schematic representing integration of sc/snRNA-seq data with ST data for cell state deconvolution. **B,C)** Cell state abundances in WTA and sc/snRNA-seq data. **B)** Healthy and COVID-19 cell state gene expression signatures were mapped to healthy and COVID-19 ROIs, respectively. Scatter plot shows the percentage of estimated cell abundance enriched in ROIs of COVID-19 (red) or healthy (blue) samples. Student’s t-test with the Bonferroni adjustment for multiple comparisons was used (****P < 0.0001). **C)** Beeswarm plots illustrate enrichment (red) or decrease (blue) of neighbourhoods in COVID-19 for each indicated cell type calculated using MiloR (FDR < 0.05). **D,E)** The distribution of selected healthy (**D**) and COVID-19 (**E**) cell state signatures across DAD pathologies. Dot plots show cell abundance values that were z-score normalised per cell type across rows/pathologies (colour) and the fraction of ROIs with cell abundance above the average value of all ROIs in a given pathology (size).

Differential gene expression (DGE) analysis of EDAD, MDAD and ODAD compared to PRES histopathological states revealed a trend of largely downregulated markers, including the shared downregulation of 371 genes, as well as 34 shared upregulated genes (Extended Fig. 5, Table S9-12). EdgeR pseudobulk analysis of EDAD vs ODAD ROIs further highlighted inflammatory and fibrotic signatures within COVID-19 tissue (Fig. 3A). Within EDAD, we observed upregulation of interleukin-related genes *IL1A*, *IL1B* and *IL6*, interferon alpha and gamma-related *IFNG*, *IFIT1,* and *MX1*, and proliferation related G2/M checkpoint markers *CDK1*, *CDC6,* and *CDC45* (Fig. 3B-C). Furthermore, EDAD enriched for *SPP1,* encoding the matricellular protein OPN, involved in leukocyte recruitment and immune cell activation (Kahles, Findeisen and Bruemmer, 2014), was previously shown to be increased in macrophages of patients with idiopathic pulmonary fibrosis (IPF) (Morse *et al.*, 2019; Hatipoglu *et al.*, 2021). In contrast, ODAD ROIs enriched for genes associated with extracellular matrix (ECM) turnover, including matrix metalloproteinases (*MMP2*, *MMP9,* and *MMP14*), as well as collagen deposition (*COL5A1*, *COL6A1*, and *COL6A2*), the pathological fibroblast marker *CTHRC1* (Tsukui *et al.*, 2020), and TGFbeta related genes (*SMAD6, SMAD7,* and *ID1*) (Fig. 3B-C). This signature infers activation of pro-fibrotic and ECM remodelling pathways in ODAD. Taken together, these data present novel biomarkers to stratify DAD stages and highlight molecular pathways underlying the progression of an inflammatory phenotype in EDAD to a profibrotic phenotype observed histologically in ODAD.

**Figure 5.**
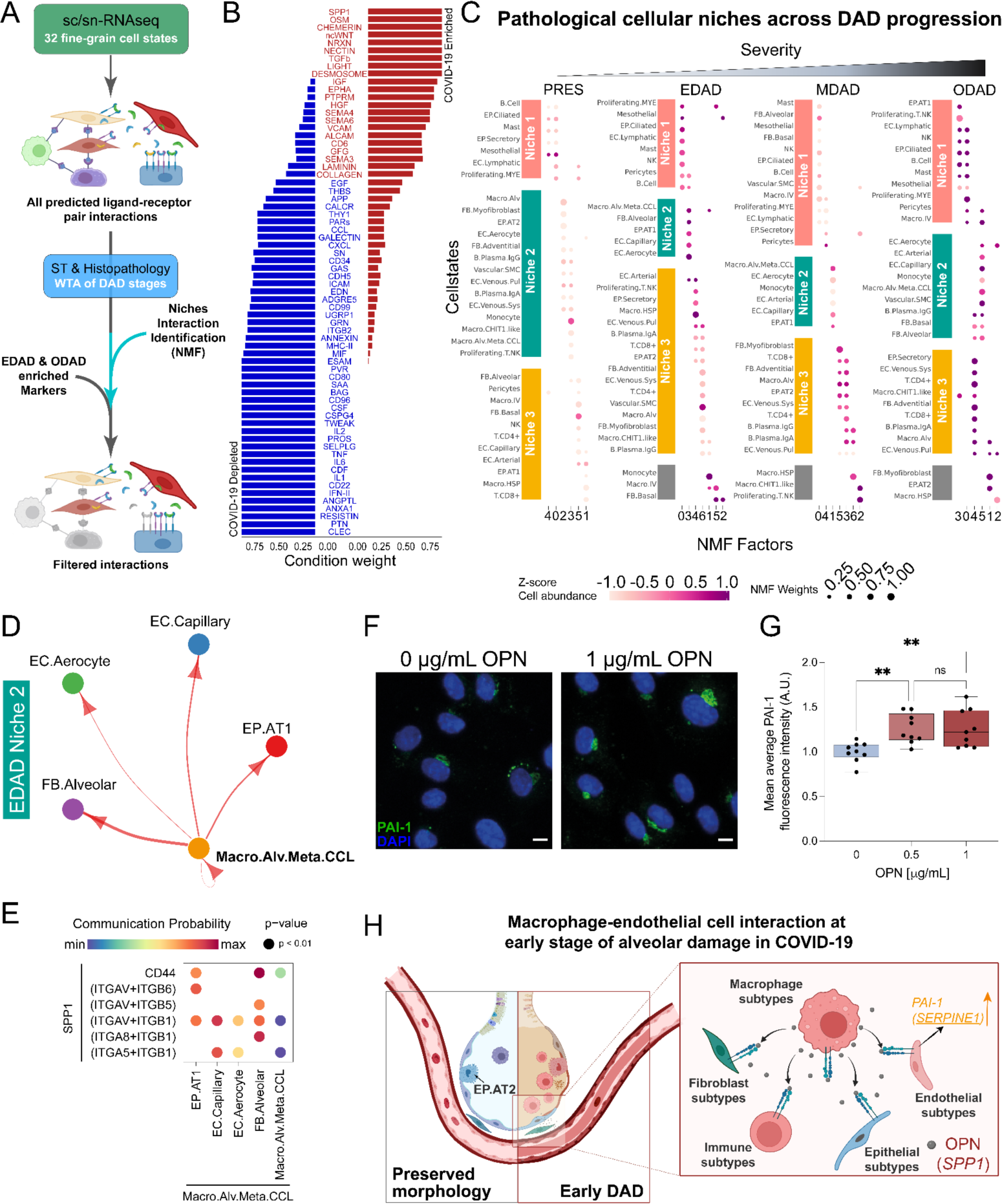
Cellular niches and cell-cell signalling across DAD stages. **A)** Diagram of analysis approach for mapping cell-cell communication across early versus late DAD. **B**) Waterfall plot visualisation of global pathway analysis between healthy (blue) and COVID-19 (red) sc/snRNA-seq, with significant pathways highlighted in blue and red respectively. **C)** NMF analysis across histopathological states. **D)** SPP1 signalling within the COVID-19 sc/snRNA-seq compartment, mapped to EDAD niche 2. Arrows indicate directionality of ligand (SPP1) signalling to receptors between cell states. **E)** Dotplot visualisation of SPP1 signalling to specific receptors across EDAD niche 2 cell states. **F)** Representative confocal images of endothelial cells treated with 1 µg/mL rhOPN or vehicle (0 µg/mL) for 24 hours. Scale bar = 10 µm. **G)** Box-whiskers plot of PAI-1 average fluorescence intensity. Values normalised to vehicle control (0 µg/mL rhOPN). n=3 independent experiments, performed in triplicate. One-way ANOVA; **p<0.01. **H)** Summary of Macrophage subtypes contributing to pro-thrombotic and anti-fibrinolytic states in early DAD through SPP1 signalling.

### Dysregulation of the coagulation cascade in COVID-19

With coagulopathy being a major complication in severe COVID-19 (Conway *et al.*, 2022; Spyropoulos and Bonaca, 2022), and frequent presence of microthrombi in the lung microvasculature (Hanley *et al.*, 2020), we interrogated the expression of genes encoding factors contributing to clot formation and resolution. DGE analysis of the sc/snRNA-seq dataset revealed differential regulation of several genes directly and indirectly involved in the fibrinolysis pathway, including upregulation of *SERPINE1,* encoding the fibrinolysis inhibitor PAI-1, in arterial, capillary, and pulmonary venous EC (Fig. 3C, Table S5). Concomitant downregulation of *PLAT*, the transcript for the fibrinolytic factor tPA, which is inhibited by *SERPINE1*/PAI-1, likely contributes to enhancing anti-fibrinolytic effects (Fig. 3D, Table S5). We also observed downregulation of *ANXA2* (Annexin A2), which typically acts as a receptor for *PLAT/*tPA to promote fibrinolysis (Valls *et al.*, 2021), as well as downregulation of *PROCR* (protein C receptor) and *THBD* (thrombomodulin), which together are required for activation of protein C, and downstream inhibition of *SERPINE1*/PAI-1 (Barranco-Medina *et al.*, 2017) (Fig. 3C-D). Other differentially regulated coagulation signatures included downregulation of anticoagulant *ANXA5* (Annexin A5), and upregulation of procoagulant *ANO6* (TMEM16F) (Fig. 3C), which inhibit and promote exposure of phosphatidylserine residues in the phospholipid bilayer, respectively, which in turn enable prothrombin complex formation (Reddy and Rand, 2020; Arndt *et al.*, 2022). We also observe downregulation of *CAV1* (Caveolin-1), which regulates the anticoagulant activity of TFPI (Tissue Factor Pathway Inhibitor), EDN1 (Endothelin-1) shown to promote Tissue Factor production (Kambas *et al.*, 2011), and *SERPING1* (Complement 1 inhibitor) which has both pro- and anticoagulant properties (Pryzdial, Leatherdale and Conway, 2022). Furthermore, while aerocytes are defined transcriptionally by an absence of *VWF* (Gillich *et al.*, 2020; Schupp *et al.*, 2021), we observe their upregulation of *VWF* in COVID-19, suggesting a phenotypic shift in the disease.

We confirmed the upregulation of *SERPINE1* in COVID-19 by single molecule fluorescence in situ hybridisation (smFISH) on COVID-19 post-mortem lung parenchyma tissue (Fig. 3E), observing a several fold increase in *SERPINE1* mRNA area staining compared to healthy control tissue (Fig. 3F). Collectively, this data suggests that *SERPINE1*, in conjunction with other coagulation related factors, may act as a key player in the fibrinolytic shutdown and persistency of microthrombi observed in COVID-19 lung endothelium from early stages of alveolar damage (Milross, Majo, Pulle, *et al.*, 2022; Spyropoulos and Bonaca, 2022). Additionally, while *SERPINE1* is not differentially expressed in ODAD compared to PRES (Table S11), previous studies have demonstrated its overexpression leads to ECM accumulation, as well as fibroblast and AT2 cell senescence in a model of IPF (Adnot, Breau and Houssaini, 2020; Rana *et al.*, 2020), suggesting long term implications of *SERPINE1*/PAI-1 in fibrosis and advanced DAD states.

### Distinct cellular composition of DAD stages

To reveal the cellular composition changes in COVID-19 and across DAD stages, we computationally deconvolved cell states in our ST data via integration with sc/snRNA-seq using the cell2location-WTA model (Roberts *et al.*, 2021; Kleshchevnikov *et al.*, 2022) (Fig. 4A). Given the pervasive transcriptional changes in COVID-19 (Delorey *et al.*, 2021) (Extended Figure 1E), we derived gene expression signatures of cell states from both healthy and COVID-19 donors from our integrated sc/snRNA-seq atlas and mapped them separately in our WTA dataset (Methods). Initially, we compared the abundance of coarse-grained cell states between healthy and COVID-19 ROIs. We observed an increase of immune subtypes (monocyte/macrophages, T & NK, and mast cells) as well as a depletion of epithelial subtypes (EP.AT1, EP.AT2, EP.Ciliated) in COVID-19 tissue (Fig.4B, Extended Figure 6A). Differential abundance analysis of our sc/snRNA-seq dataset using MiloR (Dann *et al.*, 2022) similarly revealed an enrichment of immune and depletion of epithelial related subpopulations, further confirming destruction of the alveolar bed and increased inflammation observed clinically in COVID-19 patients (Erjefält *et al.*, 2022) (Fig.4C, Extended Figure 6B). Additionally, smFISH was used to confirm the loss of AT1 and AT2 cells in COVID-19 lung parenchyma as indicated by reduced expression of *AGER* and *SFTPC*, respectively (Extended Fig. 6C).

Next, we leveraged our integrated sc/snRNA-seq and ST datasets to identify cellular abundance changes across DAD stages (Fig. 4D-E, Extended Fig. 7A). First, we examined whether our analysis recovered previously known DAD associated cellular patterns (Ashwin *et al.*, 2023; Milross *et al.*, 2023). Initially, we observed that the healthy gene expression signatures of epithelial subtypes were depleted in both PRES and DAD tissue locations compared to healthy controls (Fig. 4D), consistent with our previous report on disease phenotypes preceding morphological changes associated with DAD (Milross *et al.*, 2023). Comparing the distribution of COVID-19 cell state signatures across DAD stages, we found that fibroblast subtypes (FB.Basal and FB.Alveolar) involved in normal alveolar function (Tsukui *et al.*, 2020), as well as monocytes associated with early stages of inflammation (Italiani *et al.*, 2020) were enriched in EDAD compared to PRES and ODAD (Fig. 4E, Extended Fig. 7A). Conversely, fibroblast populations (FB.Myofibroblast and FB.Adventitial) implicated in fibrosis (Kendall and Feghali-Bostwick, 2014), were more abundant in ODAD, consistent with ODAD representing a pro-fibrotic phenotype (Cardinal-Fernández *et al.*, 2017). Additionally, lymphoid populations, including T cells (T.CD4, T.CD8) and B cells (B.Plasma.IgA, B.Plasma.IgG) (Fig. 4E, Extended Fig. 7A), were enriched in ODAD, recapitulating previous reports (Erjefält *et al.*, 2022), including our previous study on this patient cohort based on proteomic profiling (Milross *et al.*, 2023). Of note, while AT1 and AT2 cells were overall depleted in COVID-19 compared to healthy lung tissue (Fig. 4C-D Extended Figure 6), we observed that their disease cell state gene expression signatures became more abundant across progressive DAD (Extended Fig. 7). In concordance with AT1/2 cell disease phenotypes becoming more prominent across DAD, we found that many genes upregulated in AT1/2 cells in COVID-19 in our sc/snRNA-seq dataset (Extended Fig. 1E-G) showed increased expression across DAD progression in our WTA data (Extended Fig. 8). These results validate our spatial transcriptomic mapping approach and ability to stratify fine-grained cell types across DAD.

Beyond previously characterised cell state changes, we observed that distinct macrophage subtypes accumulated through progressive DAD. Macro.HSP, distinguished by heat shock protein-related markers, and Macro.Alv.Meta.CCL, were enriched in EDAD compared to PRES tissue areas (Fig. 4E). The presence of these MT+ macrophages in early DAD is consistent with our biomarker and DGE analyses (Fig. 2E, 3A). These macrophage populations were slightly reduced in MDAD and ODAD, whereas alveolar macrophages (Macro.Alv) and CHIT1+ macrophages (Macro.CHIT1.like) increased in abundance (Fig. 4E). Interestingly, these four macrophage populations were absent from COVID-19 patients that passed away from acute cardiac failure, but with the exception of Macro.Alv, were present in patients with bronchopneumonia (Extended Fig. 7C), suggesting that the superimposed bacterial infection seen in the lungs of some patients with severe COVID-19 develops on the immunological background of virus-driven EDAD. Taken together, our results define specific cellular signatures of DAD stages and identify changes in macrophage subpopulations across DAD.

### Cellular niches and cell-cell signalling across DAD stages

To explore the changes in cell-cell signalling occurring across DAD, we combined sc/snRNA-seq analysis with identification of spatially co-localised cell types and differentially expressed ligand-receptor pairs across DAD stages in the WTA data (Fig. 5A). We first used CellChat (Jin *et al.*, 2021) to infer cell-cell communication across all cell states within our sc/snRNA-seq dataset and identify differentially regulated pathways between COVID-19 and healthy control lung samples (Fig. 5B). Broadly, we found enrichment of pro-inflammatory (SPP1) and pro-fibrotic (COLLAGEN, TGFβ) pathways in COVID-19 (Fig. 5B). We also observed down-regulation of EC cell-cell adhesion signalling via ESAM, potentially reflecting loss of barrier function observed within the vasculature of COVID-19 patients, as well as downregulation of IL1, IL6 and IFN-II (interferon gamma) signalling, which may reflect an end-stage phenotype in post-mortem tissues processed for sc/snRNA-seq.

Next, we mapped pathological cellular niches across DAD progression, by assigning spatially deconvolved DAD cell states in WTA data into distinct tissue microenvironments. We used non-negative matrix factorization (NMF) on cell2location estimated abundances to identify spatially co-localizing cell states across each DAD stage (i.e. cell states that recurrently co-occur in tissue ROIs across a given pathology). We identified three major pathological niches across the stages of DAD (Fig. 5C). In niche #1, we observed epithelial, mesothelial and immune cell states. In niches #2 and #3, we observed distinct patterns of myeloid-vascular cell colocalization. In niche #2, Macro.Alv.Meta.CCL was found to co-localize with both EC.Aerocyte and EC.Capillary cells, as well as with FB.Alveolar and EP.AT1 cells. While in niche #3, Macro.Alv co-localized with EC.Venous.Pul and EC.Arterial cells, alongside FB.Myofibroblast and EP.AT2 cells (Fig. 5C). Comparing the abundances of niches across DAD progression, we observed that niches #1 and #2 were established in EDAD and persisted into ODAD. In contrast, niche #3 along with a fourth distinct cell compartment, enriching for FB.Myofibroblast, were more prominent in ODAD (Fig. 5C). These findings present distinct tissue microenvironments across early and late stages of DAD.

Finally, we examined whether any COVID-19 cellular pathways were distinctly associated with early or late DAD tissue microenvironments. Initially, we selected cellular pathways enriched in COVID-19 in our sc/snRNA-seq dataset (Fig. 5B) and examined the expression pattern of their receptor and ligands in EDAD vs ODAD pathologies in ST data by pseudobulk DGE analysis (Fig. 3A, Table S9). This revealed elevated expression of the *SPP1* ligand (encoding OPN) in early DAD (Fig 3A, Extended Fig. 9), which we had previously identified as a biomarker of EDAD (Fig. 2E). SPP1 receptors and downstream interferon genes (Platanias, 2005) were also expressed in EDAD (Extended Fig. 9A). We then examined SPP1 signalling across EDAD cell states and cellular niches. *SPP1* expression was enriched in macrophage subtypes in our COVID-19 sc/snRNA-seq data, including Macro.Alv.Meta.CCL and Macro.HSP that accumulate in EDAD (Extended Fig. 9B). In EDAD tissue microenvironments, we identified different types of macrophages as a potential source of SPP1: Macro.Alv.Meta.CCL in niche #2 and Macro.Alv and Macro.HSP in niche #3 (Fig. 5D-E, Extended Fig. 9C-D). Predicted cells receiving the signals included a range of epithelial, endothelial, mesenchymal and immune populations. Furthermore, we validated enrichment of *SPP1* upregulation in EDAD (Extended Fig. 4A), and more specifically, in niches characterised by co-localisation of macrophages and endothelial populations (Extended Fig. 10) using Visium ST datasets. These observations present macrophage *SPP1* signalling as an early event during DAD progression.

Previous studies have shown that *SPP1*/OPN can induce TGF-beta expression (Kohan, Breuer and Berkman, 2012), which in turn can upregulate *SERPINE1*/PAI-1 (Walton, Johnson and Harrison, 2017). Given the enrichment of both *SPP1* and *SERPINE1* expression in EDAD (Fig. 2E) and the dysregulation of coagulation cascade in endothelial cells in COVID-19 (Fig. 3), we explored whether *SPP1*/OPN can upregulate *SERPINE1*/PAI-1 expression in vasculature *in vitro*. We first validated our sc/snRNA-seq data by confirming the expression of *SPP1*/OPN receptors integrin alphaV-beta1 and alpha5-beta1 in cultured endothelial cells using immunofluorescence staining (Extended Fig. 9E). We then treated endothelial cells with 0.5 and 1 µg/mL rhOPN for 24hrs and observed upregulation of PAI-1 (Fig. 5F-G). Taken together, we demonstrate that macrophage derived *SPP1*/OPN may be acting at the intersection of pro-inflammatory and anti-fibrinolytic pathways via *SERPINE1*/PAI-1 upregulation in early stages of DAD (Fig. 5G-H).

## Discussion

One of the critical questions in understanding COVID-19 pathology is the advancing transcriptional regulation of early to later stage DAD. In this study, we have generated the most comprehensive single cell and spatial transcriptomic study of lung pathology in severe COVID-19 patients to date, providing the first characterisation of the cell states, tissue microenvironments, and cellular interactions that underlie different stages of DAD. By utilising a multiomics approach guided by histological analysis, these data begin to highlight the intricate gene expression changes in COVID-19 lung across various histopathological microenvironments which underlie coagulation, inflammation, and fibrosis, highlighting novel biomarkers for predicting disease severity and for therapeutic targeting. Our study provides a unique, open access resource comprising sc/snRNA-seq from COVID-19 and donor cohorts, as well as whole transcriptome spatial RNA-seq data across histopathological states of DAD.

Further, we demonstrate upregulated *SERPINE1* as a potential key regulator of the hypercoagulable state / fibrinolytic shutdown exhibited by COVID-19 patients in our sc/snRNA-seq DGE analysis (Fig. 3C). DGE analysis of the WTA data demonstrates *SERPINE1* upregulation across EDAD histopathological regions (Fig. 3A), suggesting a response in the acute phase of disease. A similar result was obtained using a random forest classifier trained on WTA ST data to predict markers of disease severity (Fig. 2E). Previous bulk analyses of lung tissue have revealed increased *SERPINE1*/PAI-1 expression in (D’Agnillo *et al.*, 2021), and increased PAI-1 protein levels in blood plasma (Zuo *et al.*, 2021). Interestingly, *SERPINE1* gene / PAI-1 protein upregulation has been linked to ACE2 inhibition via angiotensin-2 production (Kellici, Pilka and Bodkin, 2021), potentially suggesting a direct SARS-CoV-2 virus mediated mechanism for upregulated *SERPINE1* / PAI-1. As such, *SERPINE1* / PAI1 may represent a crucial biomarker for COVID-19 infection, and for targeting macro- and microthrombi therapeutically in specific vessel beds along the vascular axis.

Next, we looked at the changes in distribution of cell states across DAD states. Our multi-omic strategy, which integrated ST data with finely annotated cell types from sc/snRNA-seq, allowed identification of fine-grained cellular changes across DAD progression, including macrophage subtype changes from EDAD to ODAD (Fig. 4E). These data align with previous orthogonal profiling of cell types across DAD stages using proteomic approaches (Milross *et al.*, 2023). One limitation of our integrated mapping approach is that our sc-/snRNA-seq reference likely represents a mixture of DAD pathological states, which would likely dilute cell state signatures associated with each phenotype.

Lastly, we focused on the molecular signalling pathways within cellular niches of DAD stages. Cell-cell interaction analysis of sc/snRNA-seq data demonstrated *SPP1* (OPN) signalling enriched in macrophages in COVID-19 (Fig. 5B, 5D-E). Consistently, SPP1+ macrophages have been associated with severe COVID-19 in a recent human lung cell atlas integration effort (Sikkema *et al.*, 2022) as well as idiopathic pulmonary fibrosis (Morse *et al.*, 2019). Here, we leveraged our ST data to extend these findings and demonstrate that macrophage SPP1 signalling is enriched in early DAD and targets various endothelial, with predicted targets including epithelial, endothelial, mesenchymal and immune cells (Fig. 2E, 3A, 5C). Interestingly, *SPP1*/OPN targets a number of integrin subunits upstream of TGFbeta activation, potentially suggesting a role for this in activation of pro-fibrotic pathways. Our analysis also suggests a functional role for *SPP1*/OPN in thrombosis observed in COVID-19 via PAI-1 upregulation (Fig. 5F-H). Macrophage-derived SPP1 has also been implicated in various cancers (Gao *et al.*, 2022; Qi *et al.*, 2022), suggesting that greater exploration is required to understand the mechanism of action in disease contexts. Furthermore, OPN inhibition in a mouse model of IPF demonstrated decreased fibrosis, suggesting a candidate for therapeutic targeting (Hatipoglu *et al.*, 2021).

Taken together, our study provides a unique resource to investigate the cellular and molecular landscape of DAD progression within COVID-19 lung tissue at single cell and spatial resolution. Our data is available for interactive browsing and download at our webportal under https://covid19-multiomicatlas.cellgeni.sanger.ac.uk/.

## Data availability

The integrated sc/snRNA-seq atlas can be accessed on our webportal at https://covid19-multiomicatlas.cellgeni.sanger.ac.uk/. The spatial WTA data will be made available under the same portal. All datasets will be uploaded to ENA before full publication.

## Code availability

The custom code used for analysis of ST data and creating the figures are available on Github at https://github.com/thjimmylee/UKCIC-COVID19-paper-figures. Other code is available upon request.

## Acknowledgements

We thank the donors and their families for donating tissue samples and enabling this research. We thank Parisa Amjadi and Dominic Smith from the Xu lab (Imperial College London) for helping to establish COVID-19 sample processing under CL3 conditions, and for assistance in FACS sorting single nuclei samples. We thank Liz Tuck, Grant Calder and Tarry Porter for supporting histology and generation of WTA data,and Kristina Sorg, Erica Pawlak and Stijn van Dorgen for supporting WTA data processing. We thank L. Lawrence from the Research Histology Facility at the National Heart and Lung Institute of Imperial College London. We thank Kerstin Meyer, Ana-Maria Cujba, Amanda Oliver (Wellcome Sanger Institute), as well as Gisli Jenkins and Alison Johns (Imperial College London) for valuable discussions. We thank Jessica Cox and Martin Prete for establishing the sc/snRNA-seq and ST data portal and website. This work was made possible by funding from the UKRI (MRC) and DHSC (NIHR) for the UK CIC consortium award referenced MR/V027638/1. Additionally, this research was funded in whole, or in part, by the Wellcome Trust Grants WT206194 and 220540/Z/20/A funding to O.A.B., and a National Heart and Lung Institute PhD studentship to S.N.B. The Imperial College Healthcare NHS Trust Tissue Bank is funded by the National Institute for Health Research (NIHR) Biomedical Research Centre.

## Author Contributions

J.T.H.L., S.N.B., A.J.F, M.H., M.N. and O.A.B. conceived the study. S.N.B., P.C., B.H., M.O., A.F., P.M.K., A.J.F. and M.N. procured human lung tissue samples. S.N.B., and P.C., generated snRNA-seq data. L.M., J.M., A.F., P.M.K., and A.J.F. developed the DAD profiling strategy and identified histopathological DAD stages on lung tissue samples. V.A. and J.M. contributed to pathological assessment of lung tissue samples. K.R. and H.A. generated Nanostring WTA data. K.R. generated the smFISH data. S.N.B., and Z.J. performed the SPP1 in vitro experiments. J.T.H.L. and S.N.B. led sc/snRNA-seq and ST data analysis. J.W.C, A.A. and A.H., A.M.A.M., M.L. contributed to sc/snRNA-seq and ST data analysis. S.N.B., T.L., A.M.A.M. analysed smFISH data. B.W. and V.U. performed image analysis of WTA data. X-N.X., G.R.M., S.T., A.M.R., A.F., P.M.K., A.J.F., M.H., M.N. and O.A.B. provided project supervision and data interpretation. J.T.H.L., S.N.B., A.J.F, M.H., M.N. and O.A.B. wrote the manuscript with feedback from all authors.

## Corresponding Authors

Further information and requests for resources should be directed to Andrew Fisher, Martin Hemberg, Michela Noseda and Omer Ali Bayraktar.

## Supplementary data

**Supplementary table 1:** COVID-19 patient metadata used for snRNA-seq / RNAscope.

**Supplementary table 2:** Datasets used for generation of COVID-19 sc-/snRNA-seq object.

**Supplementary table 3:** Cell state abbreviations.

**Supplementary table 4:** Cell state differential gene expression analysis.

**Supplementary table 5:** EdgeR Pseudobulk analysis of sc-/snRNA-seq data (COVID-19 vs. Healthy / cell state).

**Supplementary table 6:** MSigDB Pathway enrichment analysis in EP.AT1 (COVID-19 vs. Healthy / cell state).

**Supplementary table 7:** MSigDB Pathway enrichment analysis in EP.AT2 (COVID-19 vs. Healthy / cell state).

**Supplementary table 8:** COVID-19 patient metadata used for WTA profiling.

**Supplementary table 9:** EdgeR Pseudobulk analysis of EDAD vs ODAD spatial WTA.

**Supplementary table 10:** EdgeR Pseudobulk analysis of EDAD vs PRES spatial WTA.

**Supplementary table 11:** EdgeR Pseudobulk analysis of ODAD vs PRES spatial WTA.

**Supplementary table 12:** EdgeR Pseudobulk analysis of MDAD vs PRES spatial WTA.

## Extended Figures

**Extended Figure 1.**
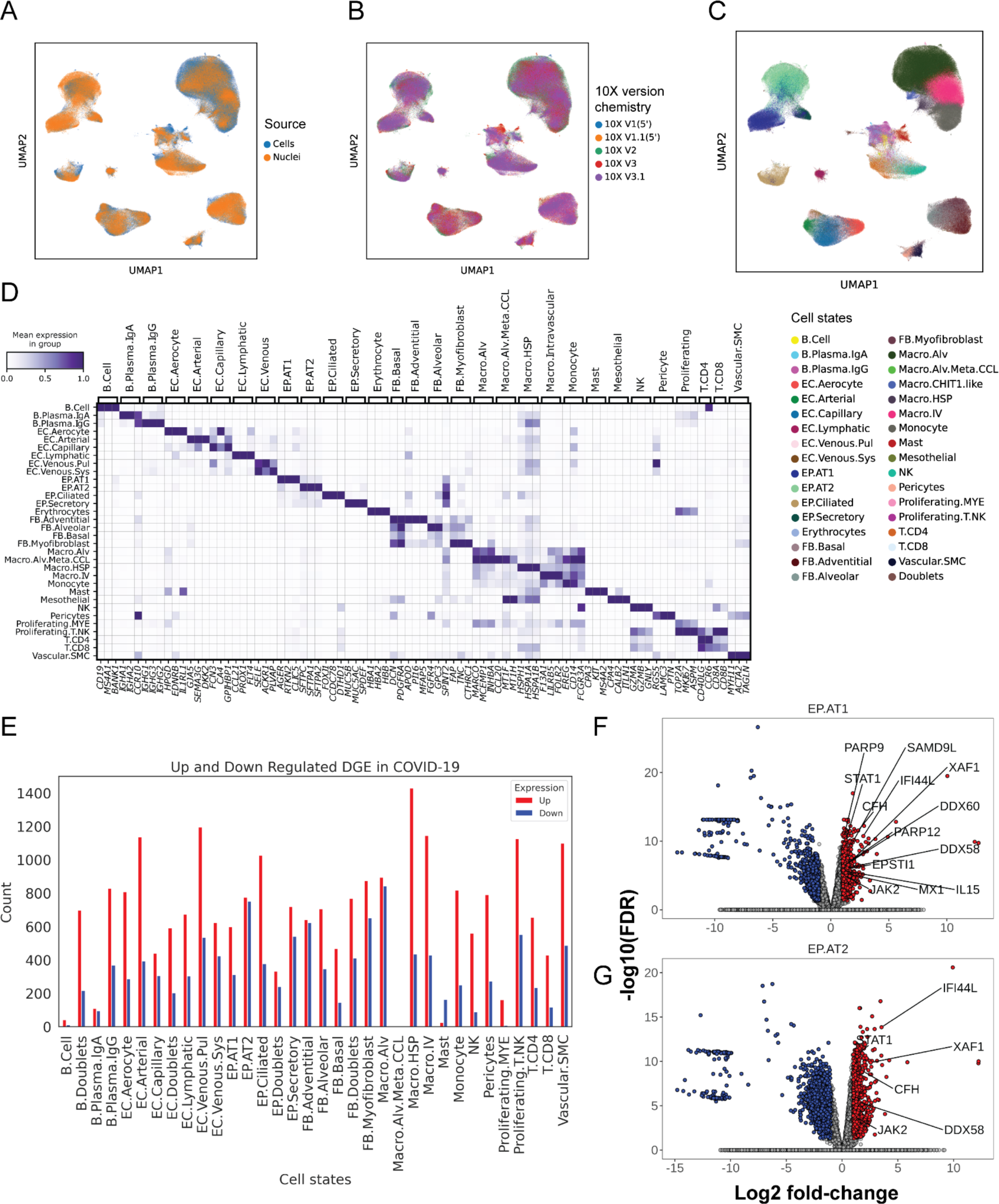
Generation of an integrated COVID19-donor lung object. **A)** UMAP representation of integrated cell or nuclei and **B)** 10X version chemistry. **C)** UMAP representation of cell states identified in COVID19-donor lung object. **D)** Matrix plot representation of cell state markers. **E)** Barplot illustrating the number of up and downregulated genes in COVID-19 per cell state (|log2fc| >1 and FDR < 0.05). Volcano plots illustrating differential gene analysis comparing COVID-19 and healthy control in **F)** EP.AT1 and **G)** EP.AT2 cell states using EdgeR with FDR=0.05 and Log2FC above 1. Red = upregulated in DADs and blue = downregulated in COVID-19.

**Extended Figure 2.**
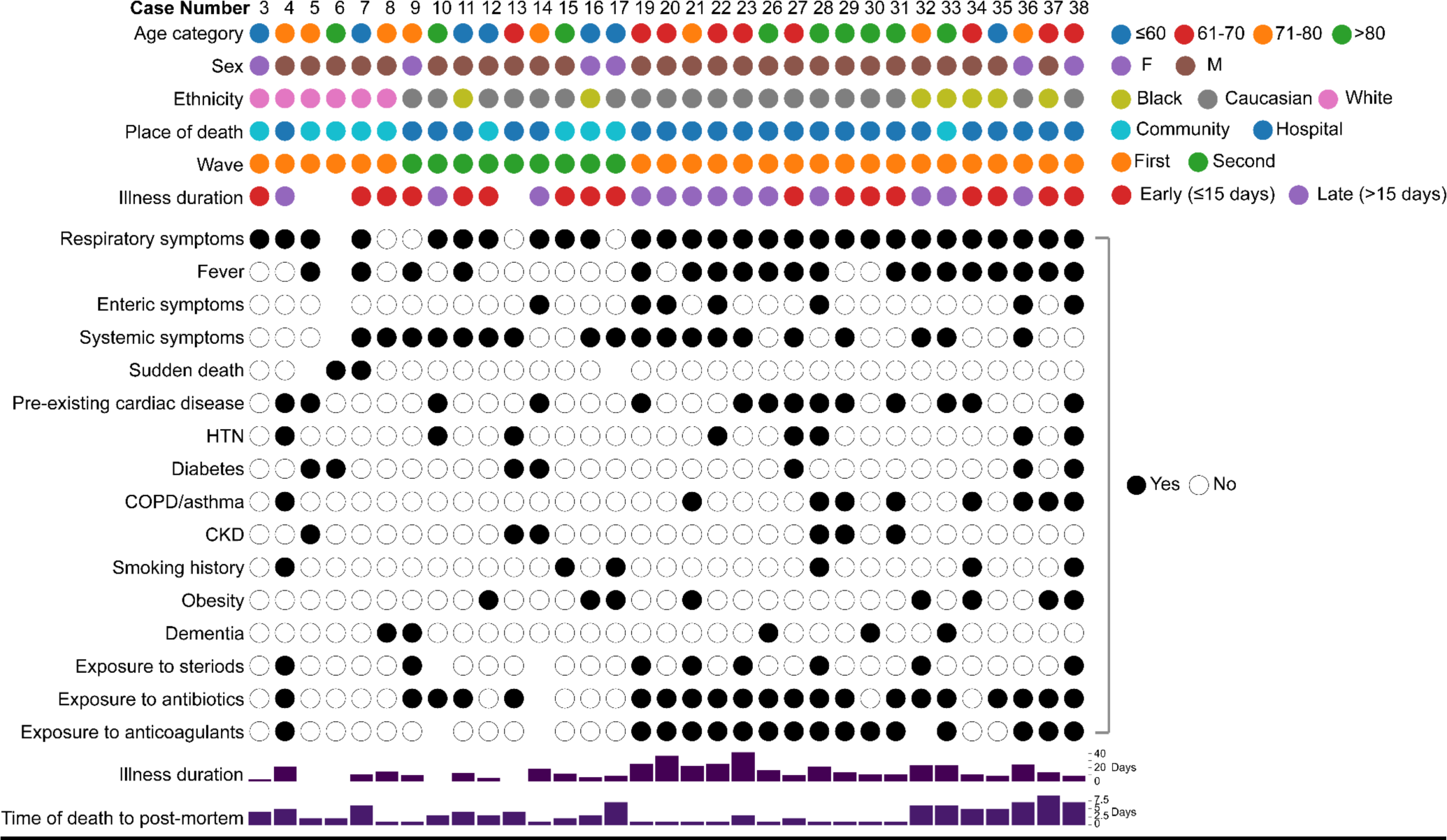
Summary of patient metadata for the WTA dataset.

**Extended Figure 3.**
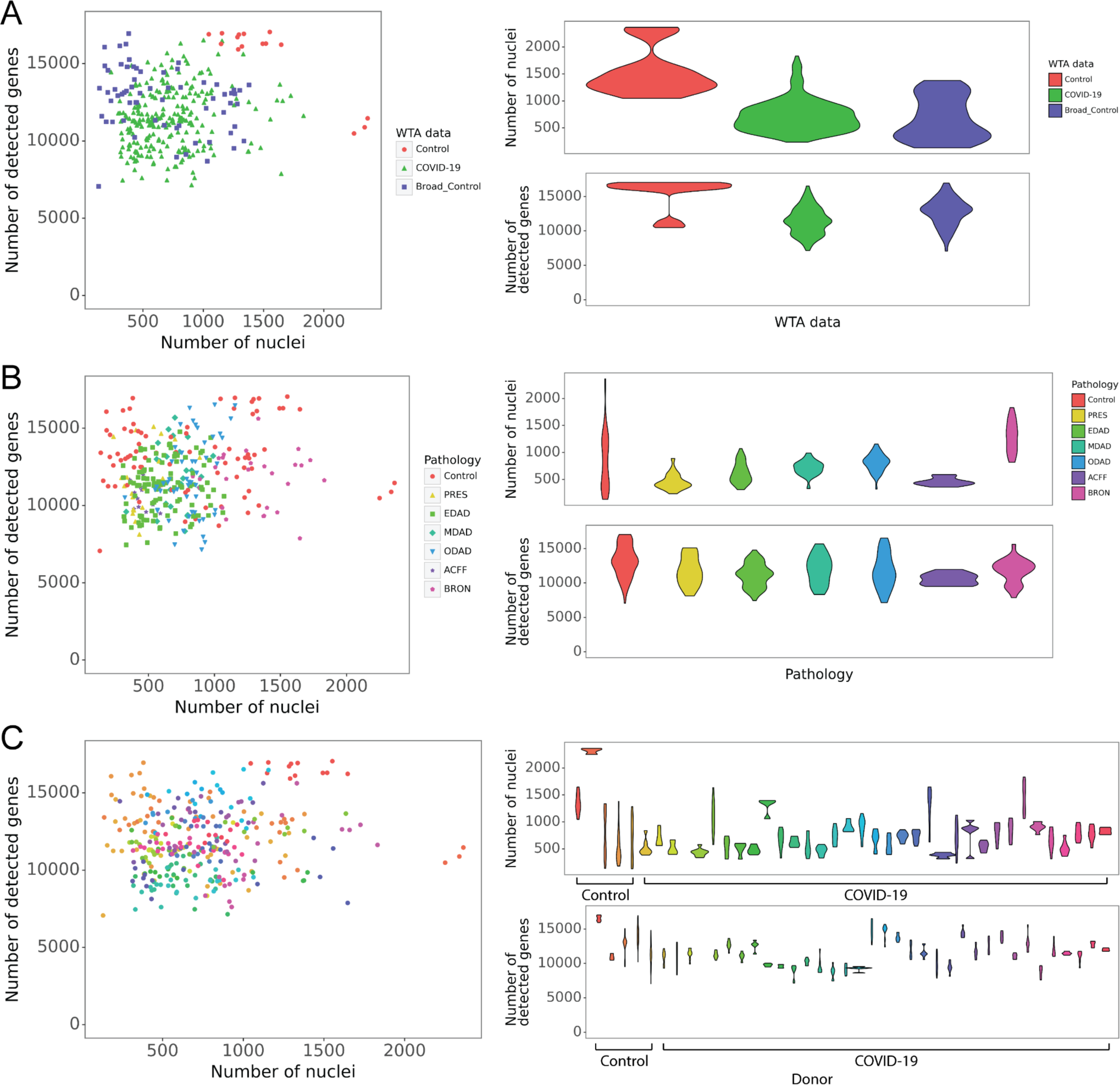
Quality control of WTA datasets. Number of nuclei (i.e. segmented from SYTO13 stains) versus number of detected genes of each ROI coloured by **A)** our WTA data from control and COVID-19 donors and the control dataset from (Delorey et al., 2021), **B)** pathologies and **C)** donors.

**Extended Figure 4.**
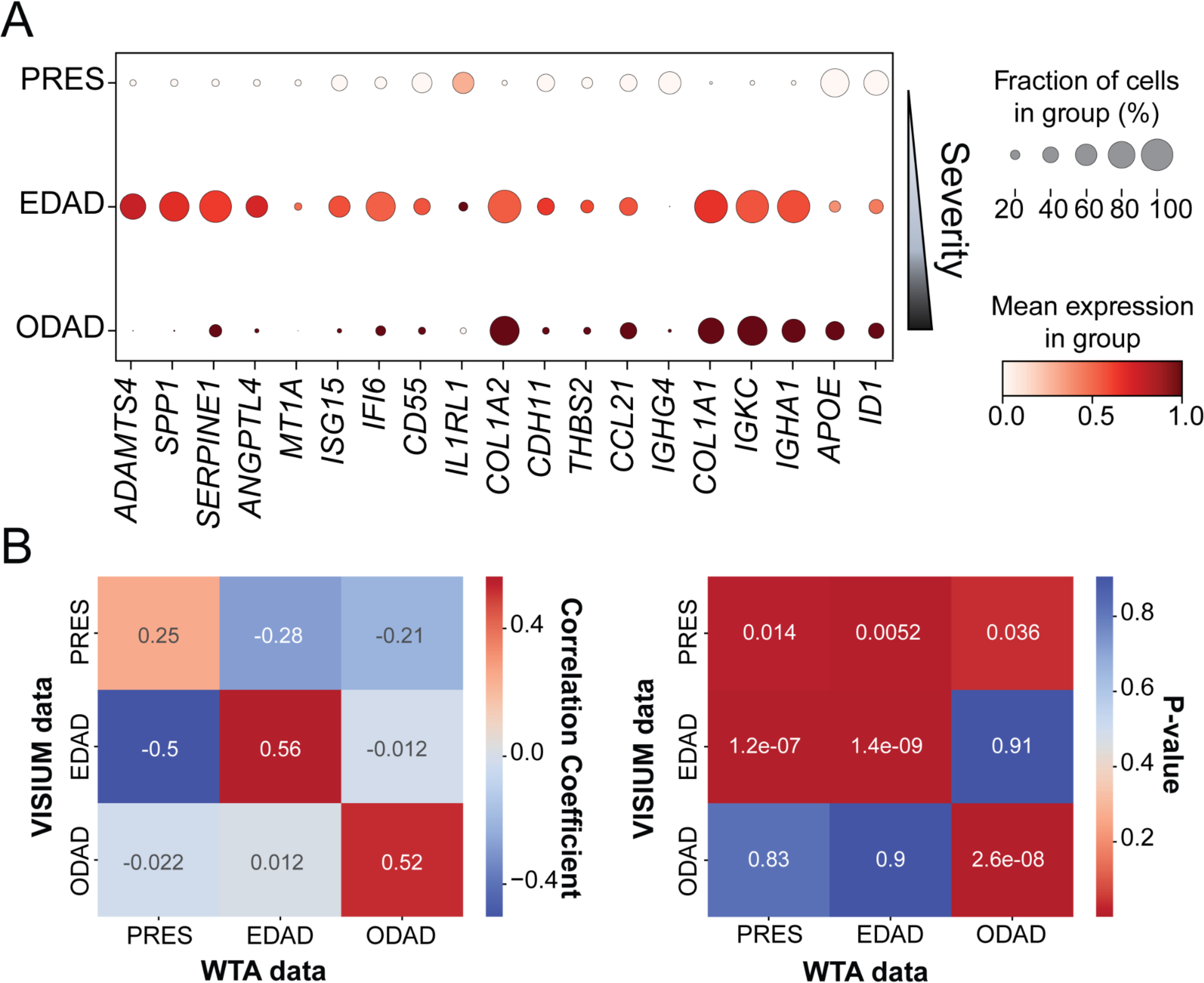
DAD biomarkers and transcriptome profiles in the orthogonal Visium dataset. **A)** Dotplot representation of histopathological state associated gene signatures obtained for random forest classifier analysis of WTA data on Visium DAD annotated regions. Dot colour = scaled mean gene expression within a group. Dot size = percentage expressed in group. **B)** Heatmaps illustrating the correlation coefficient and p-value of the transcriptome comparison between WTA data and VISIUM data of corresponding patients.

**Extended Figure 5.**
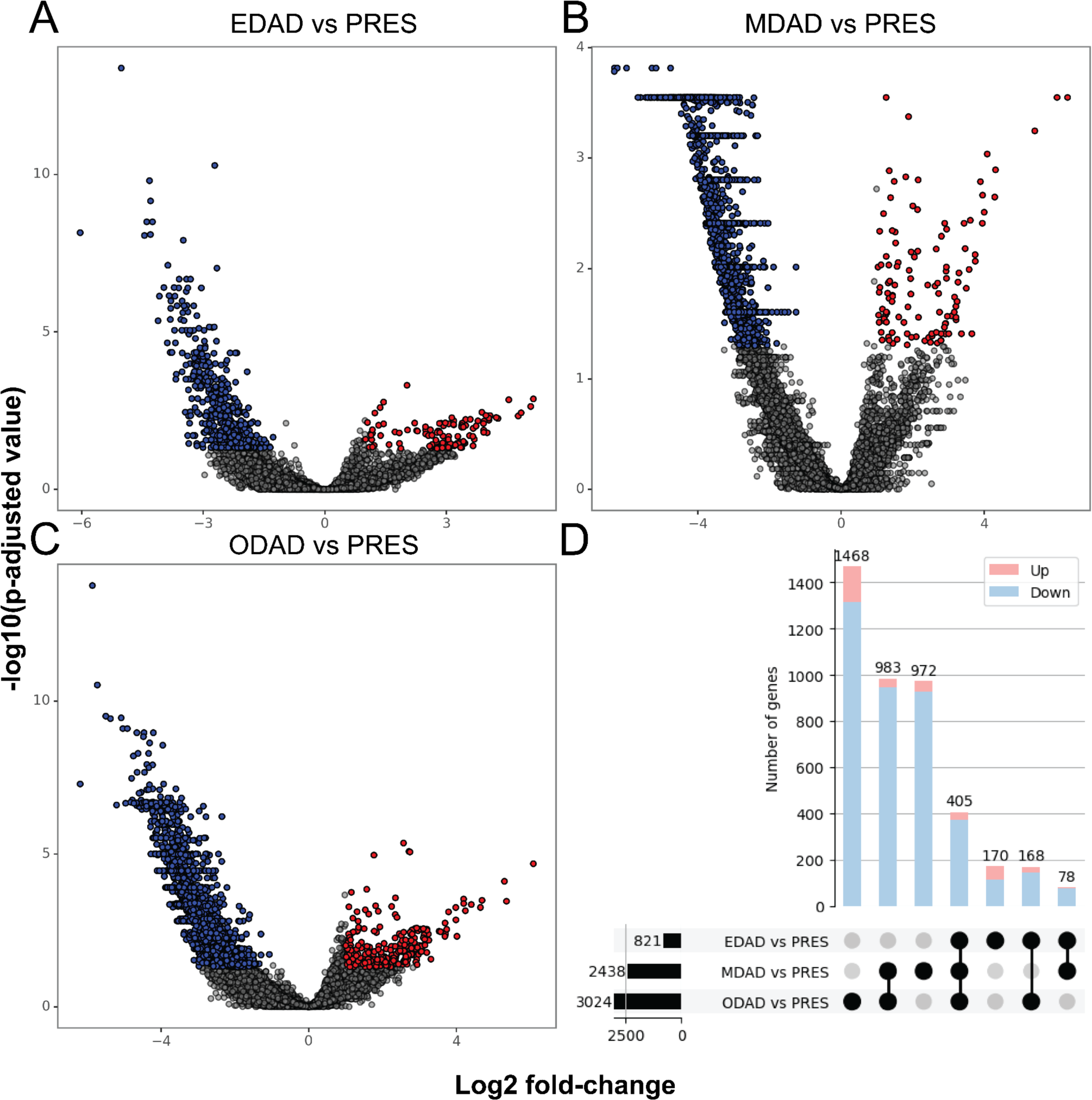
Differentially expressed genes between DAD stages and PRES tissue locations. Volcano plots illustrating differential gene analysis comparing ROIs of **A)** EDAD, **B)** MDAD and **C)** ODAD with PRES using EdgeR with FDR=0.05 and Log2FC above 1. Red = upregulated in DADs and blue = downregulated in DADs. **D)** Upset plot illustrating the intersecting DGE between EDAD vs PRES, MDAD vs PRES and ODAD vs PRES.

**Extended Figure 6.**
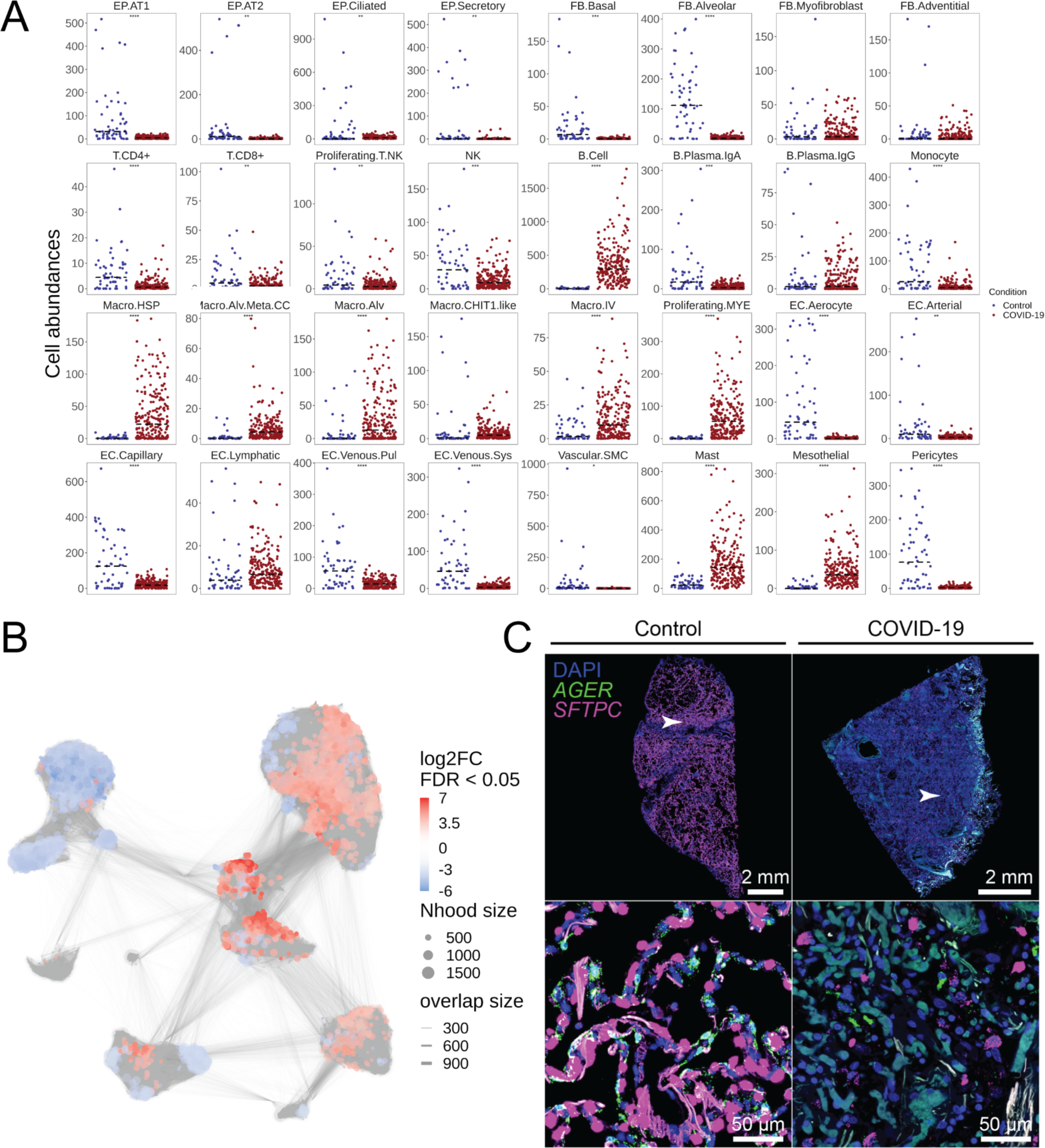
Cell abundance changes in COVID-19 samples. **A)** Spatial WTA abundance analysis between healthy control and COVID-19 samples, which were deconvolved using healthy and COVID-19 cell state gene expression signatures, respectively. Student’s t-test with the Bonferroni adjustment for multiple comparisons was used (****P < 0.0001, ***P < 0.001, **P < 0.01, *P < 0.05) **B)** UMAP representation of sc/snRNA-seq neighborhoods identified by MiloR (FDR 5%). **C)** smFISH for AGER (green), and SFTPC (magenta) counterstained with DAPI (blue) in healthy control and COVID-19 lung parenchyma samples.

**Extended Figure 7.**
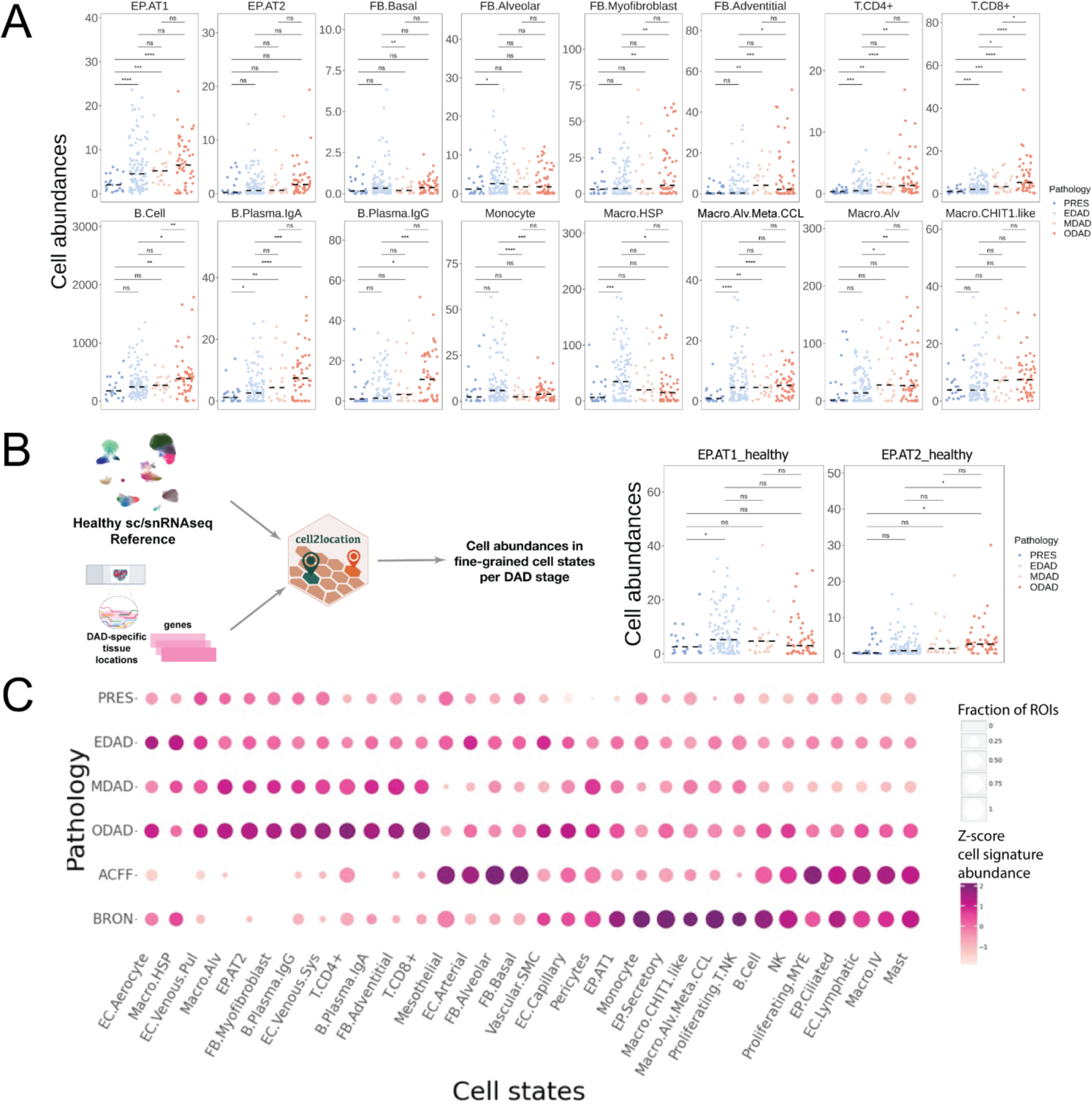
Cell abundance changes across DAD stages. **A,B)** Spatial WTA abundance analysis between DAD stages using the (**A**) COVID-19 and (**B**) healthy control cell state gene expression signatures. Student’s t-test with the Bonferroni adjustment for multiple comparisons was used (****P < 0.0001, ***P < 0.001, **P < 0.01, *P < 0.05). Median of cell abundance is indicated by the dashed line. **C)** Cellular compositions across pathologies. Shown as a dot plot of the z-score normalised by cell state (colour) and percentage of ROIs above the mean of cell abundance of cell state (size).

**Extended Figure 8.**
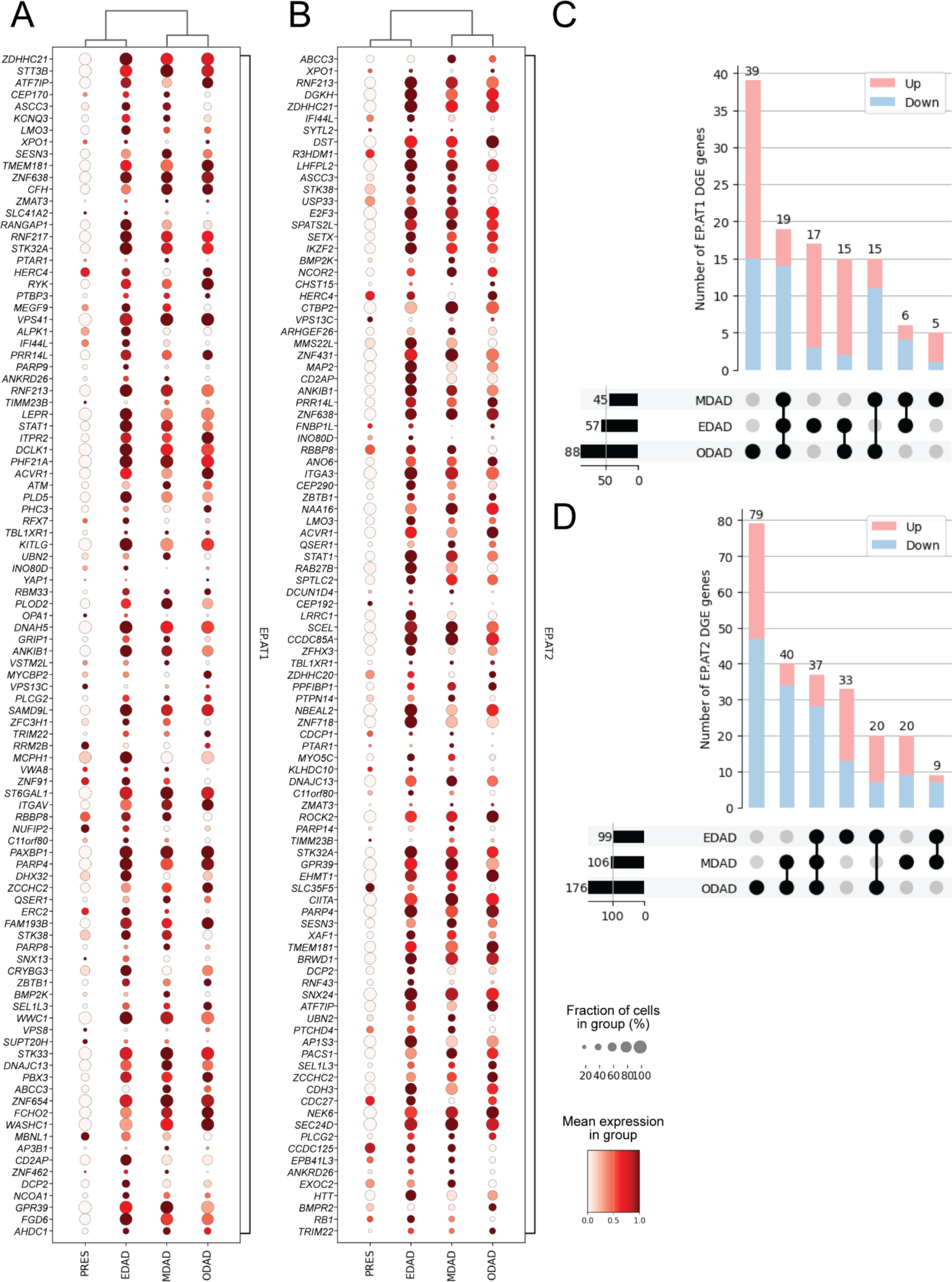
Expression patterns of AT1 and AT2 dysregulated genes across DAD stages. Genes that were dysregulated in AT1 and AT2 cells in COVID-19 were initially identified from our sc/snRNA-seq datasets using DEG analysis, then queried on our spatial WTA data of DAD stages. Dotplot visualisation of top 100 differential genes resulting from the comparison of COVID-19 and healthy control sc/snRNA-seq in **A)** EP.AT1 and **B)** EP.AT2 cells across DAD stages in the WTA dataset. Upset plots illustrating the numbers of up and down regulated DEGs in **C)** EP.AT1 and **D)** EP.AT2 cells across DAD stages.

**Extended Figure 9.**
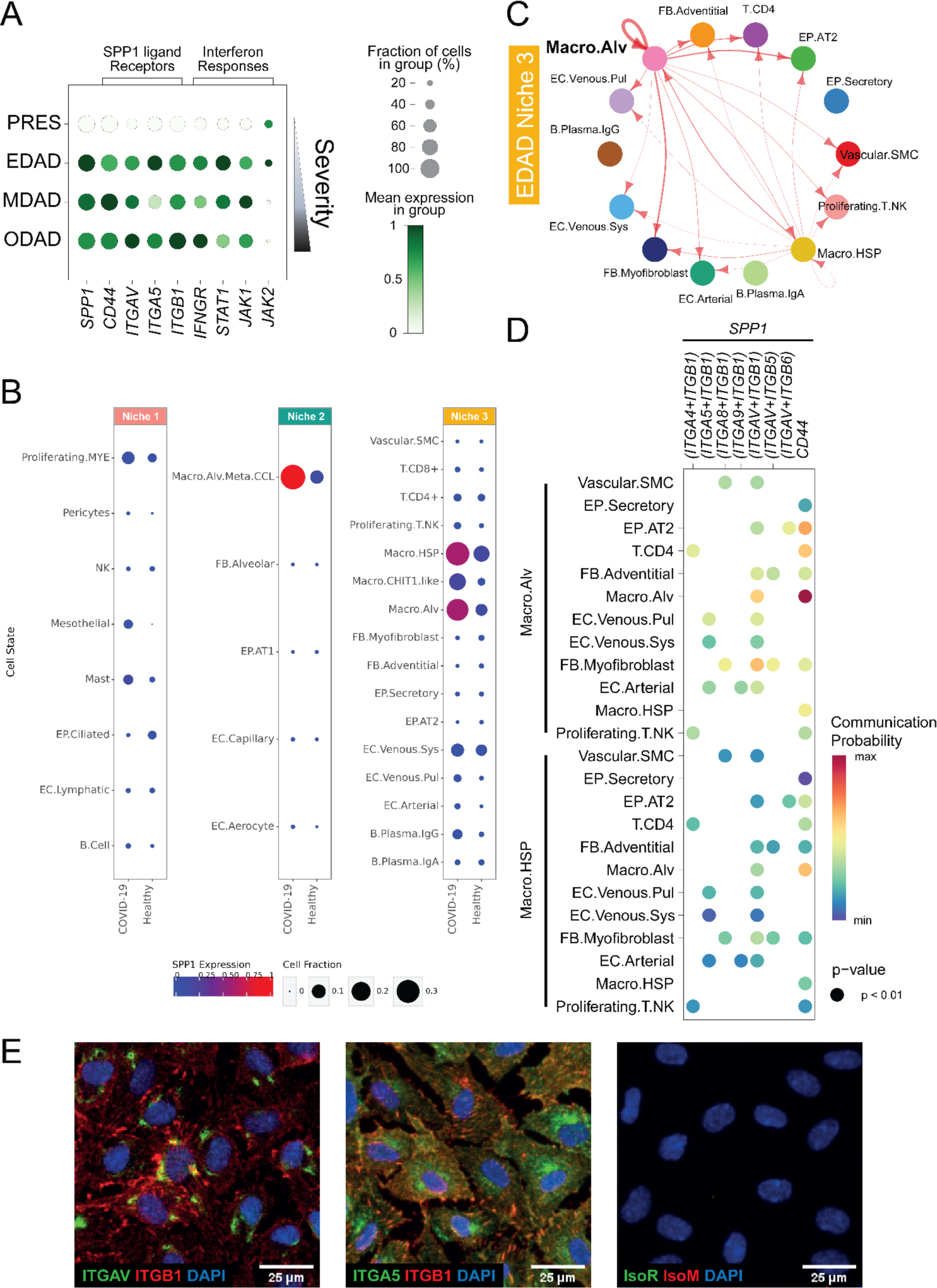
SPP1/OPN signalling in early DAD. **A)** Dotplot visualisation of SPP1 ligand and receptor expression, and interferon response associated genes across histopathological states in WTA data. **B)** sc/snRNA-seq expression pattern of SPP1 in COVID-19 and healthy control cell states of sc/snRNA-seq mapped to EDAD niches. **C)** Visualisation of SPP1 signalling within the COVID-19 sc/snRNA-seq compartment, mapped to EDAD niche 3. **D)** Dotplot visualisation of SPP1 signalling to specific receptors across EDAD niche 3 cell states. **E)** Immunofluorescent staining for OPN receptors ITGAV / ITGB1 and ITGA5 / ITGB1, as well as matched isotype controls in cultured endothelial cells (HUVEC).

**Extended Figure 10.**
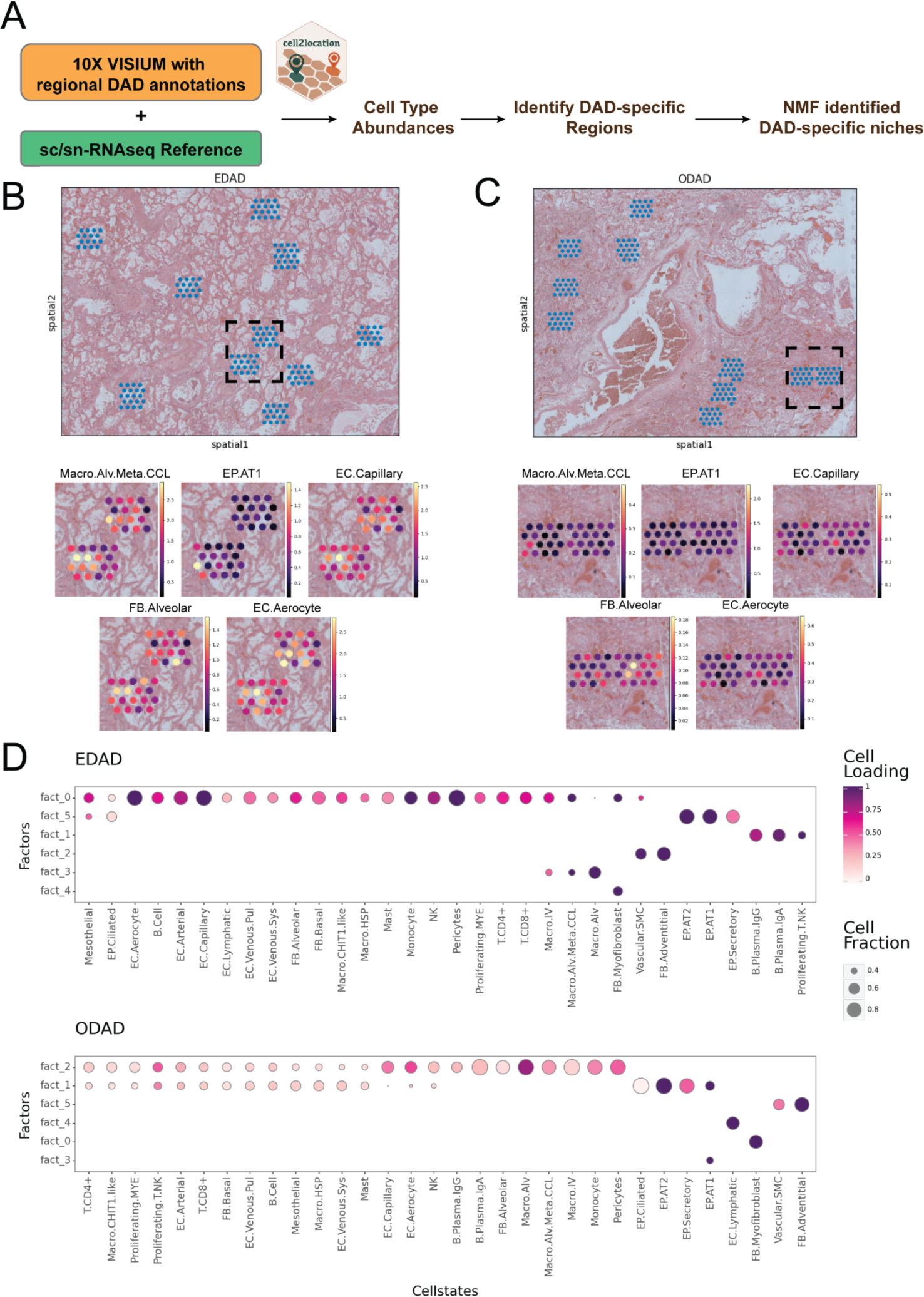
DAD-specific cellular niches in Visium data. **A)** Schematic representing integration of sc/snRNA-seq data with 10X Visium data using cell2location for downstream abundance and niche analyses. **B)** Estimated cell abundance of selected cell types within EDAD specific regions that are associated with niche 2 cellular compartments in WTA analysis. The colour scale is representing cell abundance at the 99.2% quantile **C)** Estimated cell abundance of selected cell types within ODAD specific regions that are associated with niche 2 cellular compartments in WTA analysis. The colour scale is representing cell abundance at the 99.2% quantile **D)** Identification of cell compartments using NMF. Shown as a dot plot of the estimated NMF weights (colour) and mean of cell fractions (size) of cell types (columns) across NMF.

## Methods

### Human lung tissue procurement and ethics

Human lung samples from patients who died with severe COVID-19 were obtained from 3 UK based biobanks. The Newcastle Hospitals CEPA Biobank (NHS ethics 17/NE/0070). The Imperial College Healthcare Tissue Bank (ICHTB), supported by the National Institute for Health Research (NIHR) Biomedical Research Centre based at Imperial College Healthcare NHS Trust and Imperial College London (NHS ethics 22/WA/0214). The ICECAP tissue bank at the University of Edinburgh (NHS ethics16/ED/0084). Work on these samples at the University of York was approved by the Hull York Medical School Ethics Committee (20/52). Additional samples from control donors were obtained from Cambridge Biorepository for Translational Medicine CBTM), Addenbrooke’s Hospital, Cambridge, under approval from the East of England - Cambridge South National Research Ethics Service Committee (REC 15/EE/0152). The collection of clinical metadata is described in detail in our previous studies (Ashwin *et al.*, 2023; Milross *et al.*, 2023) and the metadata is presented in Table S8.

### Tissue processing

Fresh human post-mortem tissue samples were either snap frozen in liquid nitrogen and stored at - 80°C for single nuclei transcriptomics, or fixed in 10% neutral-buffered formalin for 24-72 hours before transfer to 70% ethanol and processed to paraffin (FFPE) for WTA profiling.

### Single nuclei extraction and library preparation from COVID-19 lung post-mortem tissue

Snap-frozen COVID-19 patient post-mortem lung was mechanically dissociated using a pre-cooled pestle and mortar. Dissociated tissues were transferred to a pre-cooled Dounce homogeniser containing Homogenisation Buffer (HB) (250mM Sucrose (Sigma Aldrich, Cat#S0389); 25mM KCl (ThermoFisher, Cat#AM9640G); 5mM MgCl2 (ThermoFisher, Cat#AM9530G); 10mM Tris Buffer pH 8 (ThermoFisher, Cat#AM9855G); 1μM DTT (ThermoFisher, Cat#P2325); 1xEDTA-Free Protease Inhibitor (Sigma Aldrich, Cat#11873580001); 0.4U/μL RNAseIn Plus (Promega, Cat#N2611); 0.2U/μL SUPERasein (ThermoFisher, Cat#AM2696); 0.1% Triton X-100 (Sigma Aldrich, Cat#T8787) and Nuclease Free Water (Sigma Aldrich, Cat#W4502). Samples were further dissociated with 20 strokes of a loose and tight pestle. Cell suspensions were filtered through a 40μm strainer (Corning, Cat#352340). Nuclei suspensions were centrifuged at 500rcf for 5mins at 4°C. The supernatant was aspirated, and the pellet resuspended Storage Buffer (DPBS-/- (Gibco, Cat#14190094); 4% Bovine Serum Albumin (Sigma Aldrich, Cat#A3059); 0.2U/μL Protector RNAseIn (Sigma Aldrich, Cat#03335402001). Nuclei were stained using NucBlueTM Hoechst 33342 (ThermoFisher, Cat#R37605) and incubated for 10mins on ice. Nuclei were sorted by FACS (FACSAria^TM^III, BecktonDickinson), gated on size and DNA containing events (NucBlue^TM^ positive in the 405-450/50 channel) and collected in fresh Storage Buffer. Nuclei were counted, centrifuged at 500rcf for 5mins at 4°C and resuspended in fresh Storage Buffer at a concentration of 1000 nuclei/μL. Single cell 3’ gene expression libraries were obtained from single nuclei using 10x Chromium Next GEM Single Cell v3.1 kit. Quality control of cDNA and libraries was performed using Agilent 2100 Bioanalyser High Sensitivity DNA analysis (Agilent. Libraries were sequenced using the NovaSeq sequencing platform, targeting 50,000 reads per nucleus.

### Lung tissue histology and pathology annotation

Formalin-fixed paraffin-embedded lung blocks were obtained from multiple lung regions from each patient and were serially cut and mounted onto slides and stained with haematoxylin and eosin (H&E). The primary slide from each block was imaged using brightfield microscopy and the images were uploaded onto an OMERO webserver to serve as a reference slide for interactive annotation. Regions of interest (ROIs) were selected by a consultant histopathologist with cardiothoracic expertise with sizes ranging from 0.25mm^2^ (500µm x 500µm) to 1mm^2^ (1000µm x 1000µm), each being the selection target for a collaborative co-application of multiple advanced pathology technologies conducted across multiple academic centres. An ROI classification framework was developed based on experience gained through an early pandemic pilot population (Milross, Majo, Pulle, *et al.*, 2022) and nomenclature reflected existing published literature (Mauad *et al.*, 2021; Erjefält *et al.*, 2022). ROI classifications included the temporal phases of DAD - exudative DAD (EDAD), organising DAD (ODAD) and mixed (or ‘intermediate’) DAD (‘MDAD’) - as well as bronchopneumonia (‘BRON’) and pulmonary oedema consistent with acute cardiac failure (‘ACFF’). Detailed histological selection criteria and methodology is outlined further in Milross et al. (Milross *et al.*, 2023).

### Nanostring GeoMX Whole Transcriptome Atlas slide preparation

In addition to the use of RNase-free reagents, surfaces, equipment, and staining containers were cleaned using RNase AWAY Surface Decontaminant (Thermo) throughout slide processing. Following RNAscope staining, slides were processed according to the NanoString GeoMX RNA assay protocol (MAN-10087-02). Sections were briefly rinsed in nuclease-free water, and then post-fixed in 10% neutral-buffered formalin for 5 minutes. Fixation was quenched by incubation in 0.1 M glycine, 0.1 M Tris, twice for 5 minutes each, followed by 5 minutes washing in PBS, whereafter probes were applied immediately, again according to the assay protocol. The Whole Transcriptome Atlas (WTA) probe reagent was diluted in pre-equilibrated buffer R to a final probe concentration of 4 nM and added to each slide, which was covered with a Hybrislip cover (Grace Bio-Labs) and incubated for 18 hours at 37°C in a HybEZ II System, humidified with 2× SSC (saline-sodium citrate) buffer. The following day, slides were de-coverslipped by brief rinsing in 2× SSC with 0.05% Tween-20, and then washed twice for 25 minutes each in 2× SSC, 50% formamide at 37°C, and twice for 5 minutes each in 2× SSC at room temperature. Following washing, slides were counterstained with the DNA dye SYTO13. A SYTO13 stock (5 μM) was clarified by centrifugation at 13,000 g for 2 minutes, and then diluted to 500 nM in buffer W prior to staining in the dark for 30 minutes. Finally, slides were washed twice for 3 minutes each in 2× SSC buffer.

### Nanostring GeoMX ROI collection

Slides were covered with buffer S and loaded into the GeoMX DSP instrument for imaging and collection. Channels: SYTO13 (ex. 466-494 nm, em. 505-527 nm), CD68 (ex. 579-597 nm, em. 608-638 nm). Reference H&E images annotated by pathologists were overlaid and aligned to images. 500µm x 500µm ROIs were selected and transposed using the polygon tool. ROIs were illuminated by UV light to cleave the barcodes and the aspirate was collected. Following collection of sequencing tag-containing aspirates, wells were dried either at room temperature overnight or at 65°C for 45 mins in a heating block, and then re-suspended in 10 μl of nuclease-free water (Ambion), in order to minimise any differences due to ambient evaporation.

### Nanostring GeoMX WTA Library preparation and sequencing

ROI-derived oligos were each uniquely dual-indexed using the i5 x i7 system (Illumina). A 4 μl aliquot of each re-suspended ROI aspirate containing the photocleaved oligos was amplified in a PCR reaction containing 1 μM i5 and i7 primers and 1× NSTG PCR Master Mix. UDG digestion was carried out at 37°C for 30 min, and then deactivated at 50°C prior to denaturation at 95°C for 3 minutes, and 18 cycles of amplification: 95°C for 15 seconds, 65°C for 1 minute, 68°C for 30 seconds. Final extension was conducted at 68°C for 5 minutes.

Prior to purification, PCR products were combined into sub-pools, groups of ROIs based upon area, in order to permit biasing of sequencing to ensure sufficient coverage of smaller ROIs. Each subpool of PCR products was purified with two rounds of AMPure XP beads (Beckman Coulter) at 1.2× sample volume of beads.

Pooled libraries were quantified using an Agilent 2100 Bioanalyzer and High Sensitivity DNA Kit. The libraries were pooled in a biased manner into two sequencing reactions, each of which was sequenced with 30PE reads across all four lanes of an Illumina NovaSeq 6000 S4 flow cell at a concentration of 400 pM, with 5% PhiX spike-in, yielding 26 billion reads.

### RNAscope in situ hybridisation and immunohistochemistry

FFPE sections for both RNAscope and NanoString WTA were cut at a thickness of 5 μm using a microtome, placed onto SuperFrost Plus slides (VWR), and baked overnight at 55°C to dry and ensure adhesion. Tissue sections were then processed using a Leica BOND RX to automate staining with the RNAscope Multiplex Fluorescent Reagent Kit v2 Assay (Advanced Cell Diagnostics, Bio-Techne), according to the manufacturers’ instructions. Automated processing included baking at 60°C for 30 minutes and dewaxing, as well as heat-induced epitope retrieval at 95°C for 30 minutes in buffer ER2 and digestion with Protease III for 15 minutes. For visualisation of markers prior to NanoString GeoMX profiling, 3-plex RNAscope was developed using tyramide signal amplification with Opal 570, Opal 620, and Opal 650 dyes (Akoya Biosciences). No nuclear stain was applied at this stage. For validation staining, 3-plex or 4-plex RNAscope stains were developed using Opal 520, Opal 570, and Opal 650 dyes (Akoya Biosciences), as well as TSA-biotin and streptavidin-conjugated Atto 425 (Sigma). Nuclei were counterstained with DAPI at 167 ng/ml.

### Confocal Imaging

Imaging of validation RNAscope-stained slides was performed using a Perkin Elmer Opera Phenix High-Content Screening System, in confocal mode with 1 μm z-step size, using 20× (NA 0.16, 0.299 μm/pixel) or 40× (NA 1.1, 0.149 μm/pixel) water-immersion objectives. Channels: DAPI (excitation 375 nm, emission 435-480 nm), Atto 425 (ex. 425 nm, em. 463-501 nm), Opal 520 (ex. 488 nm, em. 500-550 nm), Opal 570 (ex. 561 nm, em. 70-630 nm), Opal 650 (ex. 640 nm, em. 650-760 nm).

### RNAscope quantification

Quantification of *SERPINE1* in COVID-19 compared to donor tissues was performed using ImageJ. Channels were normalised by subtracting the raw image with a Gaussian blur transformation with sigma=5. The value for area/n_cells was calculated by dividing the number of cells (DAPI positive nuclei) and the *SERPINE1* stain area in each sample. Donor control area/n_cell values were averaged, and used as a benchmark for calculating the log2fc of individual COVID-19 samples.

### sc/snRNA-seq integration and QC

Library samples prepared in this study were aligned to reference genome GRCh38-3.0.0 using CellRanger version 3.1.0. Publicly available data were downloaded from relevant sources (Table S2). Seurat v4 reciprocal PCA (RPCA) was applied to study batches as previously documented, with SCTransform used to correct for single cell or single nuclei as source per study batch, regressing out percent mitochondrial and ribosomal genes (Hao *et al.*, 2021). A second batch correction was performed on Seurat integrated counts using Harmony (Korsunsky *et al.*, 2019), correcting for single cell or single nuclei as source, donor and 10x version. Doublet removal was performed using Scrublet (Wolock, Lopez and Klein, 2019). Further QC was performed to filter genes (200 < nFeature_RNA < 7,500), counts (400 < nCount_RNA < 40,000), mitochondrial genes (for single cell: percent_mito < 20%, for single nuclei: percent_mito < 5%) and ribosomal genes (for single cell: percent_ribo < 20, for single nuclei: percent_ribo < 5%). Downstream analysis was performed in SCANPY (Wolf, Angerer and Theis, 2018). Clustering was performed on the global integrated object using the Leiden algorithm (Traag, Waltman and van Eck, 2019). Differentially expressed genes for each cluster were calculated using the Wilcoxon Rank Sum Test with Benjimini-Hochberg correction, and used to annotate major cell type populations based on known markers. Clusters with similar DGE profiles were merged.

### CellTypist Automated Annotation

sc/snRNA-seq data was annotated using three reference lung models trained on a logistic regression framework provided with the CellTypist tool, including ‘Cells_Lung_Airway’, ‘Nuclei_Lung_Airway’ (Madissoon *et al.*, 2022), and ‘Human_Lung_Atlas’ (Traag, Waltman and van Eck, 2019; Sikkema *et al.*, 2022). Cells and nuclei were subset and underwent automated annotation separately based on the ‘Cells_Lung_Airway’ and ‘Nuclei_Lung_Airway’ respectively. ‘Human_Lung_Atlas’ was applied to the combined cell/nuclei object. Briefly, an initial prediction step was performed using the ‘best_match’ approach, assigning one cell type label to each individual cell or nucleus, with highly variable genes not restricted. Next, over-clustering was performed using default parameters of the ‘majority_voting’ step to refine annotations based on unbiased Leiden clustering. Predicted annotations were used to guide manual subclustering annotation.

### Manual annotation of major cellular compartments

Major cellular compartments, including epithelial, endothelial, stromal, myeloid, T/NK cells and B cells were subset separately from the global object. For all compartments, the top 5,000 highly variable genes were calculated using the highly_variable_genes function in SCANPY (Wolf, Angerer and Theis, 2018), PCA was performed, and batch correction performed using HarmonyPy (Korsunsky *et al.*, 2019). Neighbours were calculated prior to UMAP and Leiden overclustering. DGE was calculated with the ‘rank_gene_groups’ function, using Wilcoxon Rank Sum Test with Benjamini-Hochberg correction. Manual annotation was performed guided by previous CellTypist predicted annotations in conjunction with known cell type / state markers, with merging of Leiden clusters with similar transcriptional profiles. Sankey plots were generated using PySankey.

### Pseudobulk DGE analysis

DGE analysis by EdgeR pseudobulk was performed for each cell state between COVID-19 and donor samples as previously described (Reichart *et al.*, 2022). Significant genes were filtered for |log2fc| >1 and log10 FDR < 0.05.

### Differential abundance analysis of sc/snRNA–seq data

Differential abundance analysis was performed using MiloR as previously documented at cell type level of sc/snRNA-seq data (Dann *et al.*, 2022).

### Gene ontology analysis

Gene ontology analysis was performed using the ShinyGO tool (Ge, Jung and Yao, 2020), with FDR threshold < 0.05.

### Generation of Expression Matrices from NanoString GeoMx WTA Data

DSP sequencing data were processed with the GeoMx NGS Pipeline as described previously (Roberts *et al.*, 2021). In brief, after sequencing, reads were trimmed, merged, and aligned to a list of indexing oligos to identify the source probe. The unique molecular identifier region of each read was used to remove PCR duplicates and duplicate reads, thus converting reads into digital counts. The limit of detection (LoD) in an ROI was defined based on the mean and standard deviation (s.d.) of log2-normalised negative probe counts. On the log scale the calculation is: LoD = mean + (2 × s.d.). All ROI with the number of detected genes lower than 6 × 10^3^ was filtered out.

### Count Correct of GeoMx WTA Data

Prior analysis, the counts in the target matrix were adjusted with negative probe counts using the python package ‘CountCorrect’.

### Pathological ROI classifier

To create a classifier for discriminating the 4 stages of DAD pathologies (PRES, EDAD, MDAD, and ODAD) of ROIs by gene expression profile from the GeoMx WTA data, the top 20% highly variable genes of the ‘CountCorrect’ matrix selected by the ‘FindVariableFeatures()’ function in the R package Seurat were considered for classification. In each round of classification, one gene was excluded and the prediction accuracy, i.e. the ratio of correct pathology classified, was computed. The performance of Support Vector Machine (SVM), Decision Tree, Naive Bayes Classifier, and Random Forest was compared. For validation, a 5-fold validation strategy was performed. Of all classifiers, Random Forest showed the best performance across all ROIs.

### Pseudobulk DGE analysis of GeoMx WTA Data

DGE analysis of the GeoMx WTA data was performed by EdgeR pseudobulk. The count matrix generated by ‘CountCorrect’ was used as input. Pairwise comparison between DAD stages of COVID-19 was performed. Significant genes were filtered for |log2fc| >1 and FDR < 0.05.

### Cell State Deconvolution and Abundance Estimation of GeoMx WTA Data

To perform cell state deconvolution, the python package cell2location-WTA (Roberts *et al.*, 2021; Kleshchevnikov *et al.*, 2022) was used. The integrated sc/snRNA-seq dataset was first subsetted into healthy and COVID-19 cells to estimate reference cell state gene expression signatures of the respective conditions, which were then used separately to train cell2location models. Reference cell state signatures were estimated by taking the mean of sc/snRNA-seq gene expression profiles per cell state. In the deconvolution step, the cell state signatures were used to decompose mRNA counts in WTA ROIs. The healthy control and COVID-19 WTA ROIs were decomposed using healthy and COVID-19 cell state signatures, respectively, with the exceptions of Fig. 4D and Extended Fig. 7B where the healthy signatures were mapped onto COVID-19 data. For the deconvolution of COVID-19 WTA data, processing all ROIs in a single batch produced results where most cell states were enriched in ACFF and BRON samples, whereas few were enriched in PRES samples (data not shown). To mitigate this, the ROIs were processed in two batches of “normal-like” and “altered” cellular morphologies determined by independent image analysis. Image texture feature descriptors were extracted from DAPI image channels of each ROI using Local Binary Pattern analysis (Ojala, Pietikäinen and Harwood, 1996) and embedded with UMAP over the first 50 principal components. The inspection of the UMAP showed two distinct clusters with (i) “normal-like” alveolar morphology including PRES and ACFF samples versus (ii) “altered” morphology including ODAD, MDAD and BRON samples. EDAD samples were distributed across both clusters and were assigned between them using a random forest classifier trained on PRES and ODAD to represent normal versus altered morphologies (data not shown).

Both the cell state gene expression profiles and the WTA data were subsetted to 11,101 common genes. The following parameters were used for the cell2location model:

– Training iterations: 20,000
– Learning rate: 0.001
– Prior on cells per location: Mean for each ROI was specified as the nuclei counts estimated by the Nanostring software for each ROI, based on DAPI stains on the image. Standard deviation was set to 10% of the mean (CV, representing prior strength, of 0.1).
– Prior on cell types per location: Mean of 6. Default CV of 1.
– Cell type combinations per location: Mean of 5. Default CV of 1.
– Prior on difference between technologies: Mean of 0.5. SD of 0.125. CV of 0.25 for both.

For visualisation, the abundances of cell state gene expression signatures were normalised by the surface area of ROIs. The alternative approach to normalise cell state abundances by the total number of nuclei in each ROI from nuclei segmentation yielded similar results (data not shown).

### Identifying cell state colocalization and tissue microenvironments

Absolute cell state abundance estimates obtained from cell2location were divided by pathology and input for NMF to identify spatially interlaced tissue compartments. For each pathology, NMF implemented in the python package ’scikit-learn’ was trained for a range of *R*={6,..,12}, and the decomposition into factors was chosen as a balance between capturing pathological stages, splitting known compartments and the cell state signature enriched in specific pathology in figure 4D. To identify the microenvironment niche, a NMF loading threshold of 0.25 was applied to keep cell states that have higher co-localisation likelihood in a given pathology. Microenvironment niches were identified according to the common cell state pattern shared between all 4 stages of DADs.

### Inferring cell-cell communication

Cell-cell interactions in COVID-19 and healthy control samples were determined using CellChat as previously documented (Jin *et al.*, 2021). Log-transformed, normalised gene counts were used without accounting for population size. The ‘RankNet’ function was used to generate heatmaps to compare pathway enrichment between COVID-19 and healthy control samples across all cell states. To resolve pathological state specificity, expression of ligands and receptors contributing to significantly enriched pathways were determined between EDAD and ODAD disease states. To infer likely interactions within pathological states, pathways were mapped to relevant disease state niches and visualised using dotplots (netVisual_bubble function), circle plots (aggregateNet function) and chord plots (netVisual_aggregate function).

### OPN treatment and immunofluorescent staining of endothelial cells

Human umbilical vein endothelial cells (HUVECs; Lonza, C2519A) were treated with recombinant human osteopontin (rhOPN; R&D Systems, 1433-OP-050) for 24 hours in serum and supplement-free growth medium (EGM, Lonza, CC-3162). Following the incubation, cells were fixed in 10% formalin in PBS, then either blocked and permeabilized in 4% (w/v) bovine serum albumin (BSA, Sigma Aldrich, A3059) and 0.2% (v/v) TritonX-100 (Thermo Fisher, 85111) in PBS (Gibco, 20012027) (PAI-1 staining) or blocked in 1% (w/v) BSA and 5% (v/v) normal goat serum (EDM Millipore, S26-100ML) in PBS (ITGAV, ITGB1, ITGA5 staining) for 30 mins at RT. Incubation with primary antibodies diluted in BSA/Triton/PBS (Rabbit anti-PAI-1) or BSA/goat serum/PBS (Rabbit anti-ITGAV, Mouse anti-ITGB1, Rabbit anti-ITGA5) was performed overnight at 4°C. Isotype controls and secondary antibody only stainings were performed as negative controls. Cells were then stained with secondary antibodies (anti-Rabbit 488, anti-Mouse 647) diluted in BSA/Triton/PBS or BSA/goat serum/PBS was performed for 1 hour at RT. Finally, cell nuclei were stained with DAPI (Invitrogen, D1306) for 15 minutes at RT. High-throughput image acquisition was carried out using Cellomics ArrayScan VTI platform (ThermoFisher), using the HCS Studio with Cellomics Scan Version 6.4.4 software (ThermoFisher). The automated Zeiss Observer Z1 epifluorescence microscope was used to acquire 12 fields per well at 10x magnification. Fluorescence intensity was recorded in channels 1-3, using the filter sets XF93 Hoechst (DAPI), XF93 FITC (Alexa488), and XF93 Cy5 (Alexa647). Confocal imaging acquisition was performed with a Zeiss LSM-780 inverted microscope, using the EC Plan Neofluar 40x/1.3 oil objective at the Imperial College London Hammersmith FILM facility using 405 nm, 488 nm, and 633 nm lasers for excitation. Image processing was performed in FIJI (v.2.1.0).

### Antibodies

Primary antibodies used in these OPN validation experiments include Anti-PAI-1 (E3I5H) XP(R) Rabbit; Cell Signaling (49536S), Anti-Integrin alpha 5 [EPR7854] Rabbit; Abcam (ab150361), Anti-Integrin alpha V antibody [EPR16800] Rabbit; Abcam (ab179475), Anti-Integrin beta 1 antibody [12G10] Mouse; Abcam (ab30394), Normal rabbit IgG isotype; Cell Signaling (3900S), Normal mouse IgG1 isotype; Santa Cruz (sc-3877), Goat anti-rabbit Alexa Fluor 488; Cell Signaling (4410S), and Goat anti-mouse Alexa Fluor 647; Cell Signaling (4412S).

